# Efficient precise *in vivo* base editing in adult dystrophic mice

**DOI:** 10.1101/2020.06.24.169292

**Authors:** Li Xu, Chen Zhang, Haiwen Li, Peipei Wang, Yandi Gao, Peter J. Mohler, Nahush A. Mokadam, Jianjie Ma, William D. Arnold, Renzhi Han

## Abstract

**Background:** Recent advances in the base editing technology have created an exciting opportunity to precisely correct disease-causing mutations. However, the large size of base editors and their inherited off-target activities pose challenges for *in vivo* base editing. Moreover, the requirement of a protospacer adjacent motif (PAM) sequence within a suitable window near the mutation site further limits the targeting feasibility. In this work, we rationally improved the adenine base editor (ABE) to overcome these challenges and demonstrated the exceptionally high efficiency to precisely edit the Duchenne muscular dystrophy (DMD) mutation in adult mice.

**Methods:** We employed a fluorescence reporter assay to assess the feasibility of ABE to correct the dystrophin mutation in *mdx*^*4cv*^ mice. The intein protein trans-splicing (PTS) was used to split the oversized ABE into two halves for efficient packaging into adeno-associated virus 9 (AAV9). The ABE with broadened PAM recognition (ABE-NG) was rationally re-designed for improved off-target RNA editing activity and on-target DNA editing efficiency. The *mdx*^*4cv*^ mice at the 5 weeks of age receiving intramuscular or intravenous injections of AAV9 carrying the improved ABE-NG were analyzed at 10 weeks or 10 months of age. The editing outcomes were analyzed by Sanger and deep sequencing of the amplicons, immunofluorescence staining, Western blot and contractile function measurements. The off-target activities, host immune response and long-term toxicity were analyzed by deep sequencing, ELISA and serological assays, respectively.

**Results:** We showed efficient *in vitro* base correction of the dystrophin mutation carried in *mdx*^*4cv*^ mice using ABE-NG. The super-fast intein-splits of ABE-NG enabled the expression of full-length ABE-NG and efficient AAV9 packaging. We rationally improved ABE-NG with eliminated off-target RNA editing activity and minimal PAM requirement, and packaged into AAV9 (AAV9-iNG). Intramuscular and intravenous administration of AAV9-iNG resulted in dystrophin restoration and functional improvement. At 10 months after AAV9-iNG treatment, a near complete rescue of dystrophin was measured in *mdx*^*4cv*^ mouse hearts. The off-target activities remained low and no obvious toxicity was detected.

**Conclusions:** This study highlights the promise of permanent base editing using iABE-NG for the treatment of monogenic diseases, in particular, the genetic cardiomyopathies.

## Introduction

Mutations in the dystrophin gene cause Duchenne muscular dystrophy (DMD)^1^, a fatal genetic muscle disease affecting approximately 1 in ∼5000 male births worldwide^2^. Previous studies showed that exon deletion through CRISPR genome editing can restore dystrophin expression and function^3-17^. Although promising, this strategy raises potential safety concerns as it relies on repair of the double strand DNA break (DSB) created by CRISPR/Cas9^18-20^, which may cause unwanted large deletion and even DNA rearrangement^21-23^.

Through fusing the CRISPR-Cas9 nickase with nucleobase deaminases (e.g. cytidine or adenine deaminase), a new paradigm-shifting class of genome editing technology, termed “base editors”, have recently been developed^24-26^. DNA base editors, via catalyzing the conversion of one base to another, directly and precisely install point mutations into chromosomal DNA without making DSBs. Therefore, base editing can be developed as a promising therapeutic to correct the genetic diseases without DNA cleavage. In particular, the adenine base editors (ABEs) show remarkable fidelity in mouse embryos and plants as compared to cytosine base editors (CBEs)^27, 28^, making them highly attractive in therapeutic development. Moreover, nearly half of the point mutations causing human diseases are G-to-A or C-to-T, highlighting the potential of ABEs in correcting a large number of human diseases.

A recent study showed that *in vivo* base editing can correct a custom-made mouse model of DMD^29^, which carries a nonsense mutation in exon 20 with a classical 5’-TGG protospacer adjacent motif (PAM) sequence in the noncoding strand for recognition by the Cas9 from *Streptococcus pyogenes* (SpCas9). In silico analysis of the ClinVar database showed that about 42.8% of the 53469 human disease-causing mutations could be potential targets for base editing correction; however, the majority (∼72.4%) of these potential targets could not be suitable for SpCas9 base editing due to the lack of the 5’-NGG PAM sequence within the suitable distance from the mutations. Several variants of SpCas9 have recently been engineered with relaxed PAM (such as xCas9-3.7^30^, SpCas9-NG^31^ and ScCas9^32^) and non-G PAM^33, 34^. These enzymes greatly increase the target scope for correcting human mutations. However, their performance to correct genetic mutations in preclinical animal models remains to be determined. Here we explore the feasibility and long-term efficacy of correcting a commonly used mouse model of DMD, *mdx*^*4cv*^ mice^35^, using NG-targeting base editors.

## Methods

### Mice

Mice (C57BL/6J and B6Ros.Cg-*Dmd*^*mdx-4Cv*^/J) were purchased from the Jackson Laboratory and maintained at The Ohio State University Laboratory Animal Resources in accordance with animal use guidelines. All the experimental procedures were approved by the Animal Care, Use, and Review Committee of the Ohio State University.

### Plasmid construction

The pCMV-ABE7.10, pCMV-ABE-xCas9(3.7) and pCMV-ABEmax were obtained from Addgene. The NG mutations were introduced by fusion PCR of pCMV-ABEmax and subcloned into pCMV-ABEmax to make pCMV-ABEmaxNG. The A56G and V82G mutations were introduced into TadA* domain by fusion PCR and cloned into pCMV-ABEmaxNG to generate pCMV-iABEmaxNG. The CfaN minigene was synthesized by IDTdna and fused at the amino acid 573 of SpCas9-max through PCR amplification. The TadA-TadA*-SpCas9max(2-573)-CfaN fragment was PCR amplified and subcloned into pAAV under the control of meCMV promoter to generate pAAV-ABEmaxN-temp. The hU6 promoter with *mdx*^*4cv*^-targeting gRNA was PCR amplified and cloned into pAAV-ABEmaxN-temp to make pAAV-ABEmaxN. The CfaC fused with SpCas9max(574-end) was generated by PCR and cloned into pAAV-ABEmaxN-temp to make pAAV-ABEmaxC. Similarly, pAAV-ABEmaxN2 and pAAV-ABEmaxC2NG with the Gp41-1 intein were constructed. The *mdx*^*4cv*^ gRNA oligos were annealed and ligated into pLenti-ogRNA. The *mdx*^*4cv*^ reporter oligos were annealed and ligated into pLKO-puro-2A-mdx^4cv^-EGFP.

### Generation of AAV particles

AAV vectors were produced at the viral vector core of the Nationwide Children’s Hospital as previously described ^4^. The ABE-NG and gRNA targeting *mdx*^*4cv*^ mutation (GTTaTCTCCTGTTCTGCAGC <u>TGT</u>; note: the underlined PAM sequences were not included in the gRNA) expression cassettes were packaged into AAV9 capsid using the standard triple transfection protocol ^36^. A quantitative PCR-based titration method ^37^ was used to determine an encapsulated vector genome titer utilizing a Prism 7500 Fast Taqman detector system (PE Applied Biosystems Grand Island, NY USA). The following primers/probes were used: 5’-GGATTTCCAAGTCTCCACCC-3’ and 5’-TCCCACCGTACACGCCTAC-3’ for titering AAV9-NG, and AAV9-iNG was titered using digital droplet PCR. Titers are expressed as DNase resistant particles per ml (DRP/ml) and rAAV titers used for injection in mice were 8.9 × 10^12^ DRP/ml (AAV9-NG) and 3.0 × 10^13^ DRP/ml (AAV9-iNG).

### Cell culture and transfection

HEK293 cells were cultured in Dulbecco’s modified eagle’s medium (DMEM) (Corning, Manassas, VA) containing 10% fetal bovine serum (FBS) and 1% 100x penicillin– streptomycin (10,000 U/ml, Invitrogen). Cells were plated in 6-well plates and transfected with the 2 µg plasmids (0.5 µg reporter, 0.75 µg gRNA and 0.75 µg ABE) per well unless specified otherwise by polyethylenimine (PEI) as previously described ^38^.

### Flow cytometry

At 72 hours post transfection, HEK293 cells transfected with ABE plasmids were collected from 6-well plate and analyzed on Becton Dickinson LSR II (BD Biosciences) to determine GFP-positive cells. A total of 100,000 cell events were collected and data analysis was performed using the FlowJo software (Tree Star, Ashland, OR, USA).

### Intramuscular and intravenous administration of AAV9 particles

AAV9-NG viral particles (2 × 10^11^ vg, 25 µl) were injected into the right gastrocnemius compartment of the male *mdx*^*4cv*^ mice at 5-6 weeks of age. For systematic delivery, the male *mdx*^*4cv*^ mice at 5-6 weeks of age were administered with AAV9-NG, AAV9-iNG or AAV9-GFP viral particles (1 × 10^14^ vg/kg) via tail vein injection.

### Serological analysis

Blood samples were collected at various time points after intramuscular or intravenous injection. The blood samples were allowed to clot for 15 min to 30 min and centrifuged at 5000 rpm for 10 min in room temperature. The supernatant was collected as serum and stored at −80 °C for the biochemical assays. Measurement of ALT (BioVision Incorporated), AST (BioVision Incorporated), BUN (Arbor Assays, Michigan, USA) and cardiac Troponin I (Life Diagnostics, Inc) were performed according to the manufacturer’s protocols.

### Antibody ELISA

Antibodies against AAV9 and SpCas9 were detected by adapting previously published protocols ^39-41^. In brief, recombinant AAV9 (2×10^9^ vg/well) and SpCas9 protein (0.27 μg/well) were diluted in 1x Coating Buffer A (BioLegend) and used to coat a 96-well Nunc MaxiSorp plate. Proteins were incubated overnight at 4 °C to adsorb to the plate. Plates were washed four times 5 min each with PBS plus 0.05% Tween-20 and then blocked with 1x Assay Diluent A (BioLegend) for 1 h at room temperature. The anti-AAV2 (A20, cat. # 03-65155, American Research Products, Inc) and anti-SpCas9 antibody (Diagenode C15310258) was used as positive control for detection of anti-AAV9 and anti-SpCas9 antibodies, respectively. Serum samples were added in 1:50 dilution and plates were incubated for 2 h at room temperature with shaking. Plates were washed four times 5 min each and 100 μl of blocking solution containing goat anti-mouse IgG (Sigma 1:3,000) was added to each well and incubated at 1 h at room temperature. Plates were washed four times 5 min each, 100 μl of freshly mixed TMB Substrate Solution (BioLegend) was added to each well, and incubated in the dark for 20 min. The reaction was stopped by adding 100 μl 2N H_2_SO_4_ Stop Solution. Optical density at 450 nm was measured with a plate reader.

### Muscle contractility measurements

At 5 weeks after intramuscular AAV9-NG or intravenous AAV9-iNG injection, muscle contractility was measured using an *in vivo* muscle test system (AuroraScientific Inc). Mice were anesthetized with 3% (w/v) isoflurane and anesthesia was maintained by 1.5% isoflurane (w/v) during muscle contractility measurement. Maximum plantarflexion tetanic torque was measured during a train of supramaximal electric stimulations of the tibial nerve (pulse frequency 150 Hz, pulse duration 0.2 ms).

### Histopathological assessment of tissues

Mice were sacrificed at various time points, and tissues (heart, lung, diaphragm, spleen, kidney, liver, quadriceps and gastrocnemius) were harvested for histological, histochemical, biochemical and molecular analyses. For immunohistological examinations, tissues were embedded in optimal cutting temperature (OCT, Sakura Finetek, Netherlands) compound and snap-frozen in cold isopentane for cryosectioning. The tissues were stored at −80°C and processed for biochemical analysis and histology assessment. Frozen cryosections (7 µm) were fixed with 4% paraformaldehyde for 15 minutes at room temperature. After washing with PBS, the slides were blocked with 3% BSA for 1 hour. The slides were incubated with primary antibodies against dystrophin (ab15277, 1:100, Abcam) and laminin-α2 (ALX-804-190-C100, 1:100, Enzo) at 4 °C for 1 hour. After that, the slides were washed extensively with PBS and incubated with secondary antibodies (Alexa Fluor 488 goat anti-rat IgG, Invitrogen, Carlsbad, CA or Alexa Fluor 568 donkey anti-rabbit IgG, Invitrogen) for 1 hour at room temperature. The slides were sealed with VECTASHIELD Antifade Mounting Medium with DAPI (Vector Laboratory, Burlingame, CA). All images were taken under a Nikon Ti-E fluorescence microscope (magnification 200x) (Nikon, Melville, NY). Laminin-α2-positive and dystrophin-positive muscle fibers were counted using NIS-Elements AR version 4.50 (Nikon, Melville, NY). The amount of dystrophin positive muscle fibers is represented as a percentage of total laminin-α2-positive muscle fibers.

### Western blot analysis

Mouse tissues from *mdx*^*4cv*^ mice treated with or without AAV9-NG or AAV9-iNG were lysed with cold RIPA buffer supplemented with protease inhibitors and extracted protein samples were separated by SDS-PAGE (BioRad, 4-15%) and transferred onto Nitrocelluloase membranes (0.45 μm). The rabbit polyclonal anti-dystrophin (E2660, 1:500, Spring Bioscience, Pleasanton, CA), rabbit polyclonal anti-Cas9 (C15310258-100, 1:1000, Diagenode, Denville, NJ) and rabbit monoclonal anti-Gapdh (#2118, 1:2000, Cell Signaling Technology, Danvers, MA) antibodies were used for immunoblotting analysis. HRP conjugated goat anti-mouse (1:4000) and goat anti-rabbit (1:4000) secondary antibodies were obtained from Cell Signaling Technology, Danvers, MA. The membranes were developed using ECL western blotting substrate (Pierce Biotechnology, Rockford, IL) and scanned by ChemiDoc XRS+ system (BioRad, Hercules, CA). Western blots were quantified using Image Lab 6.0.1 software (Bio-Rad Laboratories, Hercules, CA) according to the manufacturer’s instruction.

### Extraction of genomic DNA and total RNA, PCR and Sanger sequencing

Genomic DNA from mouse tissues and cultured HEK293 cells were extracted using DNeasy Blood & Tissue Kit (Qiagen, Germantown, MD). Total RNA was extracted from mouse tissues and HEK293 cells using Quick-RNA MiniPrep Kit (ZYMO Research, Irvine, CA). Five μg of treated RNA was used as template for first-strand cDNA synthesis by using RevertAid RT Reverse Transcription Kit (Life Technologies, Carlsbad, CA). Aliquots of the RT product were used for RT-PCR analysis of dystrophin editing. PCR reactions were carried out with 100 ng genomic DNA or cDNA in the GoTaq Master Mix (Promega) according to the manufacturer’s instruction. The primers used for mouse dystrophin genomic DNA were: mDMD-i52-F; 5’-GAGGTAATAGAGCCAAGCCCT and mDMD-i53-R; 5’-GCAAGAATTCCACTTTTCACTTCCT. The primers used for mouse dystrophin mRNA were: mDMD-E51-F; 5’-CTGTCATCTCCAAACTAGAAATGC and mDMD-E55-R; 5’-GCAGCCTCTTGCTCACTTACTC. The PCR products were purified using the Wizard SV Gel and PCR Clean-up System (Promega). Purified genomic DNA and RT PCR products (100 ng) were subjected to Sanger sequencing with mDMD-i52-F and mDMD-E51-F, respectively, at the Genomics Shared Resource of the Ohio State University Comprehensive Cancer Center. The sequencing data were analyzed by BEAT program^42^.

### Targeted deep sequencing

The on-target and off-target loci were first amplified by genomic DNA PCR and/or RT-PCR using gene-specific primers with Illumina adapters (primers are provided in Supplementary **Table S2**). The first PCR products were purified using a commercial purification kit (Promega, Madison, WI, USA), diluted, pooled, and subjected to a second round PCR with primers including the index sequences. The final PCR products were electrophoresed on an agarose gel, showing a single sharp peak. The quality and quantity were assayed using an Agilent Bioanalyzer 2100 (Genomics Shared Resource, Ohio State University Comprehensive Cancer Center). The purified amplicons were pooled and sent for sequencing using a MiSeq nano-scale flow cell (Paired-end 300 base-pair reads) at The Genomics Services Laboratory of Nationwide Children’s Hospital. The FASTQ files were analyzed using CRISPResso2 ^43^ with default parameters.

### Statistical analysis

The data were expressed as mean ± S.E.M. and analyzed with GraphPad Prism 8.0.1 software (San Diego, USA). Statistical significance was determined using one-way ANOVA followed by Bonferroni post hoc-tests for multiple groups or student’s *t*-test for two groups. A *P* value of less than 0.05 is regarded as significant.

## Results

### *In vitro* reporter assay demonstrates the feasibility to correct a dystrophin mutation using ABE-NG

The *mdx*^*4cv*^ mouse carries a premature stop codon (CAA-to-TAA) in the exon 53 of *Dmd* gene^35^, which disrupts the expression of dystrophin and leads to the development of muscular dystrophy. Targeting the noncoding strand with ABEs could potentially correct this nonsense mutation. However, in the noncoding strand, there is a lack of 5’-NGG sequence at the downstream of this mutation within the suitable editing window, but a 5’-TGT PAM is present with the mutated A located at position 4 in the guide RNA (gRNA) (**Fig. 1a**), making it feasible to correct the stop codon with the NG-targeting base editors in this widely used mouse model of DMD. We first constructed a reporter plasmid^44^ with the targeting sequence from the *mdx*^*4cv*^ mice (**Fig. 1b**). The nonsense mutation in the *mdx*^*4cv*^ targeting sequence disrupts the expression of downstream EGFP and successful editing of the nonsense mutation is indicated by the restoration of EGFP expression. As shown in **Fig. 1c**, transfection with the reporter alone resulted in minimal background fluorescence. Similarly, co-transfection with the reporter, *mdx*^*4cv*^-gRNA and ABEmax failed to restore EGFP expression. However, ABE-NG (based on SpCas9-NG) successfully restored EGFP expression in this reporter assay. In contrast, ABE-x (based on xCas9-3.7) was found to be less efficient in restoring EGFP expression even though xCas9-3.7 was also engineered to target 5’-NG PAM, consistent with previous reports that xCas9-3.7 is generally less efficient than SpCas9-NG^31, 45^. FACS analysis showed that ABE-x and ABE-NG restored EGFP expression in 10% and 20% cells, respectively (**Fig. 1d, e**). These *in vitro* studies showed that ABE-NG could potentially correct the nonsense *mdx*^*4cv*^ mutation.

**Fig. 1:**
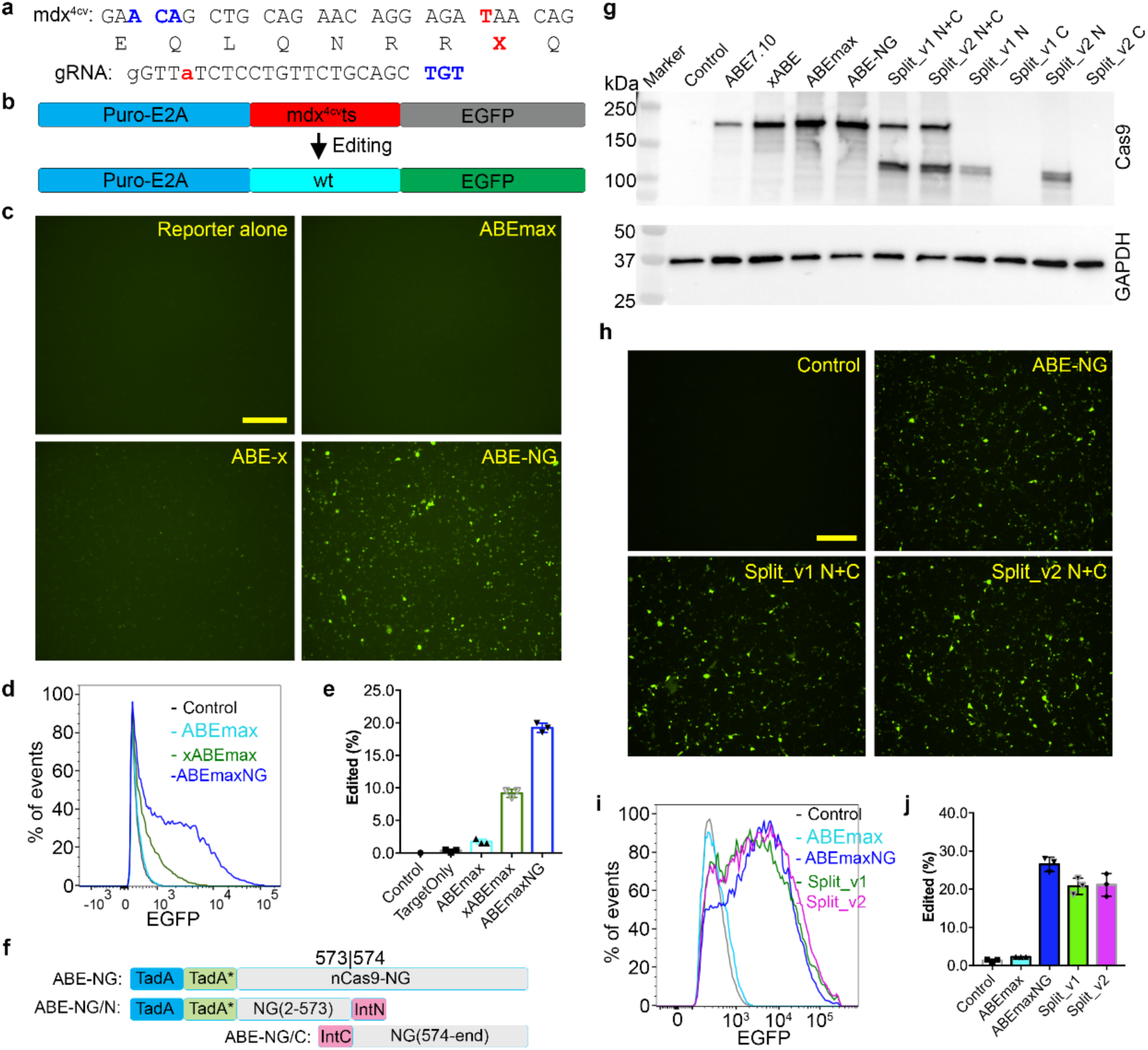
*In vitro* studies of *mdx*^*4cv*^ mutation correction using ABE-NG and its intein-split versions. **a**, Genomic DNA, encoded amino acids and guide RNA with PAM (highlighted in blue) sequences at the stop codon mutation site (red). **b**, The reporter construct contains a puromycin resistance cassette fused with E2A peptide, *mdx*^*4cv*^ target sequence and ATG-removed EGFP. Correction of the stop codon within the target sequence would allow EGFP expression. **c**, Fluorescence microscopy images of HEK293 cells transfected with reporter alone, or reporter, gRNA and one of the base editors (ABEmax, ABE-x and ABE-NG). Scale bar: 500 µm. **d-e**, Flow cytometry analysis of EGFP expression in HEK293 cells transfected as described in **c. f**, Schematics of the intein split ABE-NG. The N-terminal and C-terminal intein sequences reconstitute the full-length protein when co-expressed within the cell. **g**, Western blot analysis of HEK293 cell lysates transfected with different versions of ABEs. **h**, Fluorescence microscopy images of HEK293 cells transfected with reporter alone, or reporter, gRNA and one of the base editors (ABE-NG, split_v1 N+C or Split_v2 N+C). Scale bar: 500 µm. **i-j**, Flow cytometry analysis of EGFP expression in HEK293 cells transfected as described in **h**.

### Intein-split allows assembly of full-length ABE-NG and editing

The large size of the ABE-NG and other base editors poses a major challenge for viral packaging and *in vivo* delivery. A dual trans-splicing adeno-associated virus (AAV) approach was previously used to deliver ABE^29^ and a dual protein trans-splicing (PTS) approach using the split-intein moiety from *Nostoc punctiforme* (*Npu*)^46^ was used to deliver CBE^47^. We attempted to adopt the PTS approach to deliver ABE. In our initial preliminary experiments, the ABE was split between the ecTad-ecTadA* and the Cas9 nickase with *Npu* intein moieties, and this split renders low editing efficiency (**Supplementary Fig. S1**). To improve the editing efficiency of the split ABE, we chose the amino acid position 573 and 574 of the Cas9 nickase as the splitting site because previous studies showed that 573/574 split Cas9 exhibited near the full-length Cas9 activity^46^. Moreover, split at this site would produce roughly equal size of the two halves for AAV packaging (**Fig. 1f**). We reasoned that the split ABE could be further improved by using inteins with fast rate of PTS. We selected two inteins with remarkably fast rate of PTS: Cfa (*t*_1/2_ = 20 s at 30 °C)^48^ and gp41-1 (*t*_1/2_ = 5 s at 37 °C)^49^, which are ∼2.5-fold and ∼ 10-fold faster than the rate reported for the *Npu* DnaE intein (*t*_1/2_ = 50 s at 37 °C)^46^, respectively. Transfection of both split versions into HEK293 cells resulted in robust expression of full-length ABEs as detected by the anti-Cas9 antibody (**Fig. 1g**), although the expression level was generally lower than the ABEmax but higher than the original ABE7.10. Co-transfection with the split ABE-NG, *mdx*^*4cv*^-gRNA and the *mdx*^*4cv*^ reporter restored EGFP expression to a similar level as the full-length ABE-NG (**Fig. 1h-j**). There was no significant difference between the Cfa and gp41-1 intein splits (**Fig. 1j**). We chose the gp41-1 version for further *in vivo* experiments.

### Intramuscular delivery of AAV9-ABE-NG leads to restoration of dystrophin expression and functional improvement

We packaged the two gp41-1 intein split halves of the ABE-NG into AAV9 (**Fig. 2a**, hereafter referred to as AAV9-NG) and tested if *in vivo* delivery of ABE-NG could correct the mutation in *mdx*^*4cv*^ mice. A mini-CMV promoter with a murine creatine kinase enhancer element^50^ (meCMV, **Supplementary Fig. S2**) was used to drive the expression of two halves of ABE-NG. Each half also carries a U6-gRNA expression cassette to increase the expression of gRNA as previous studies showed that increased gRNA expression could enhance genome editing ^13, 51^. The two halves of AAV9-NG (2×10^11^ vg) were injected into the right *gastrocnemius* muscle of *mdx*^*4cv*^ mice at the age of 6 weeks, and the mice were analyzed seven weeks after injection.

**Fig. 2:**
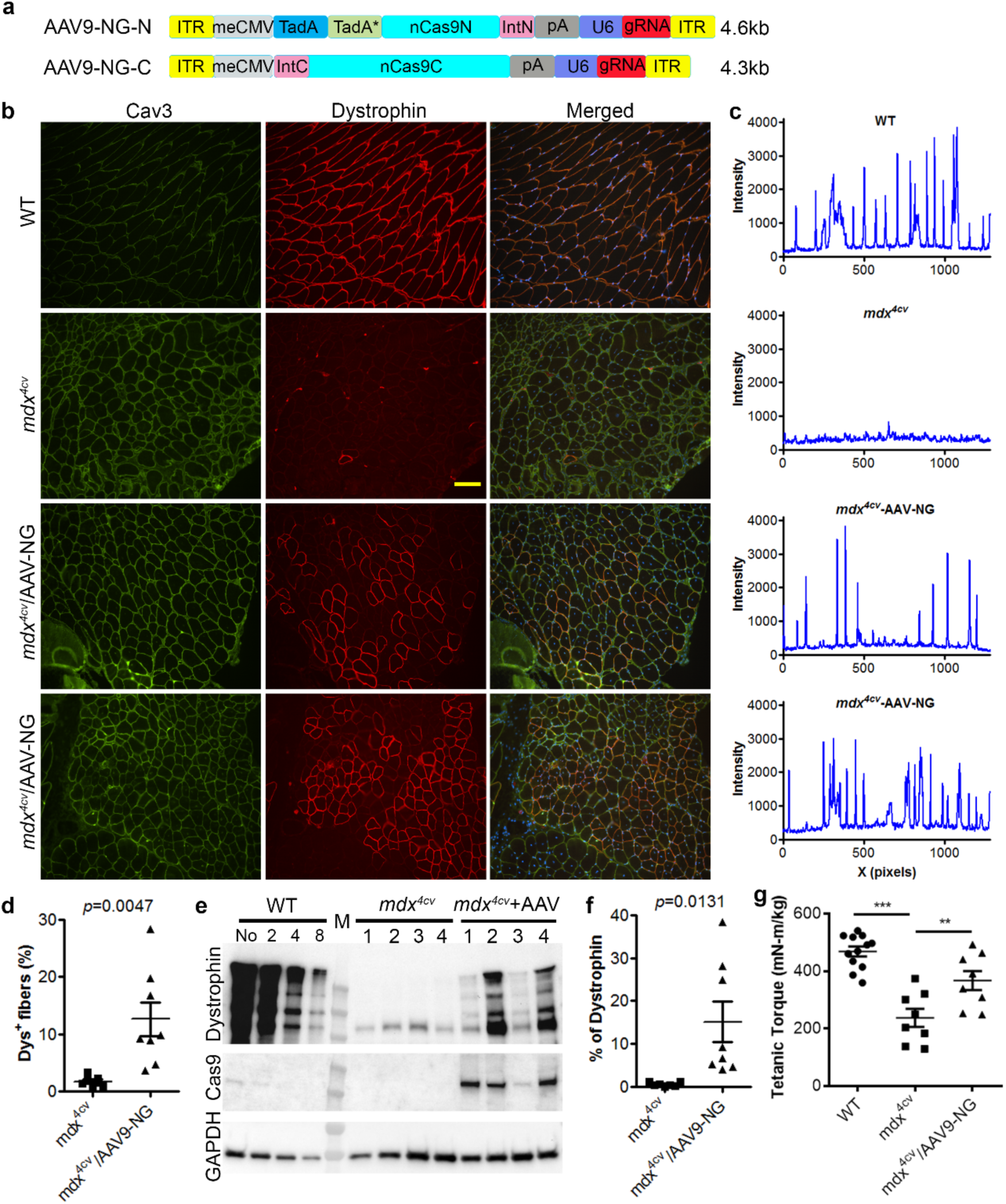
Intramuscular administration of AAV9-NG corrected *mdx*^*4cv*^ mutation and restored dystrophin expression. **a**, Schematics of the AAV9 vectors with each of the intein split ABE-NG halves and the gRNA expression cassette. **b**, Dystrophin and caveolin-3 co-immunostaining of gastrocnemius muscle sections from WT and *mdx*^*4cv*^ mice with or without intramuscular injection of AAV9-NG. Scale bar: 100 µm. **c**, Line profile of dystrophin-stained images showing the dystrophin fluorescence intensity. **d**, Quantification of dystrophin-positive (Dys^+^) fibers. **e**, Western blot analysis of muscle homogenates with anti-dystrophin, cas9 and Gapdh antibodies. The WT muscle lysates were loaded at different dilutions (No dilution, 2x, 4x, and 8x) while the *mdx*^*4cv*^ muscle lysates were loaded at 30 µg per lane. **f**, Densitometry quantification of Western blot data. **g**, Tetanic torque measurements of the posterior compartment muscles. ***p < 0.001; **p < 0.01 (one-way ANOVA).

Immunofluorescence staining showed that dystrophin was disrupted in the contralateral skeletal muscle of *mdx*^*4cv*^ mice, while in AAV9-NG treated *mdx*^*4cv*^ muscles, dystrophin was restored at the sarcolemma with an average 12.6 ± 3.0% (n=8) of dystrophin-positive fibers (**Fig. 2b, d**). The fluorescence intensities in dystrophin-positive muscle fibers in the treated *mdx*^*4cv*^ muscles were similar to those in WT animals (**Fig. 2c**), indicating that dystrophin expression level in these corrected muscle fibers was not remarkably different from WT muscle fibers. Different mice showed some variations in dystrophin-positive fibers ranging from 3.6% to 28.3%. Moreover, α-sarcoglycan and neuronal nitric oxide synthase (nNOS), which were severely reduced at the sarcolemma of *mdx*^*4cv*^ muscles, were also restored after AAV9-NG delivery (**Supplementary Fig. 3**), indicating that the entire dystrophin-glycoprotein complex (DGC) was rescued.

We further analyzed the expression of dystrophin by Western blot analysis. Similar to the immunofluorescence staining results, dystrophin was restored to different degrees ranging from 4.0% to 38.4% of the WT level on Western blot (**Fig. 2e, f**). This variation was unlikely due to low ABE-NG expression from poor injection as we observed that the muscle with high ABE-NG expression (as probed by the anti-Cas9 antibody) could have low dystrophin rescue (**Fig. 2e**). Thus, other factors such as the recently reported off-target RNA editing activity of ABEs52, 53 may be the underlying causes of the variation in the *in vivo* base editing outcomes.

The muscle tissues contain many different cell types, which limit our capacity to precisely determine the DNA editing efficiency in muscle fibers. To estimate the editing efficiency of the dystrophin gene, we extracted the total RNA from the muscle tissues treated with or without AAV9-NG, amplified the target region by RT-PCR, and analyzed the resulting amplicons by Sanger sequencing and BEAT program^42^. The AAV9-NG treated *mdx*^*4cv*^ muscles showed up to 11% T-to-C editing at the premature stop codon (**Supplementary Fig. S4**).

To test if intramuscular AAV9-NG delivery could improve muscle function, muscle contractility was examined at 8 weeks of age using an *in vivo* muscle test system^54^. Maximum plantarflexion tetanic torque was measured during supramaximal electric stimulation of the tibial nerve at 150 Hz. Compared to WT controls, *mdx*^*4cv*^ mice produced significantly reduced torque, while intramuscular AAV9-NG injection significantly improved the tetanic torque of *mdx*^*4cv*^ muscles (**Fig. 2g**).

### Rational design improves the editing efficiency and specificity of ABE-NG

We next attempted to re-design the ABE-NG in order to improve the *in vivo* base editing outcomes. First, we replaced the dimeric adenine deaminase domain (ecTadA-ecTadA*) with a monomeric ecTadA* variant (**Fig. 3a**) to eliminate the off-target RNA editing activity. A previous study showed that the WT ecTadA domain can be removed from ABE ^55^ and the off-target RNA editing activity can be eliminated by mutagenesis ^52, 53, 55^. Using the reporter assay, we confirmed that the ABE-NGm (the monomeric TadA* with A56G and V82G mutations fused with SpCas9-NG nickase) maintained the on-target DNA editing activity (**Fig. 3b**). To examine the off-target RNA A-to-I editing activities, we amplified and sequenced five RNAs which were previously shown to be highly modified by ABEmax in human cells ^55, 56^. As expected, transfection of HEK293 cells with ABE-NG induced high levels of A-to-I RNA editing in all these transcripts (**Fig. 3c**); however, such A-to-I RNA editing was essentially eliminated in cells transfected with ABE-NGm (**Fig. 3c**).

**Fig. 3.**
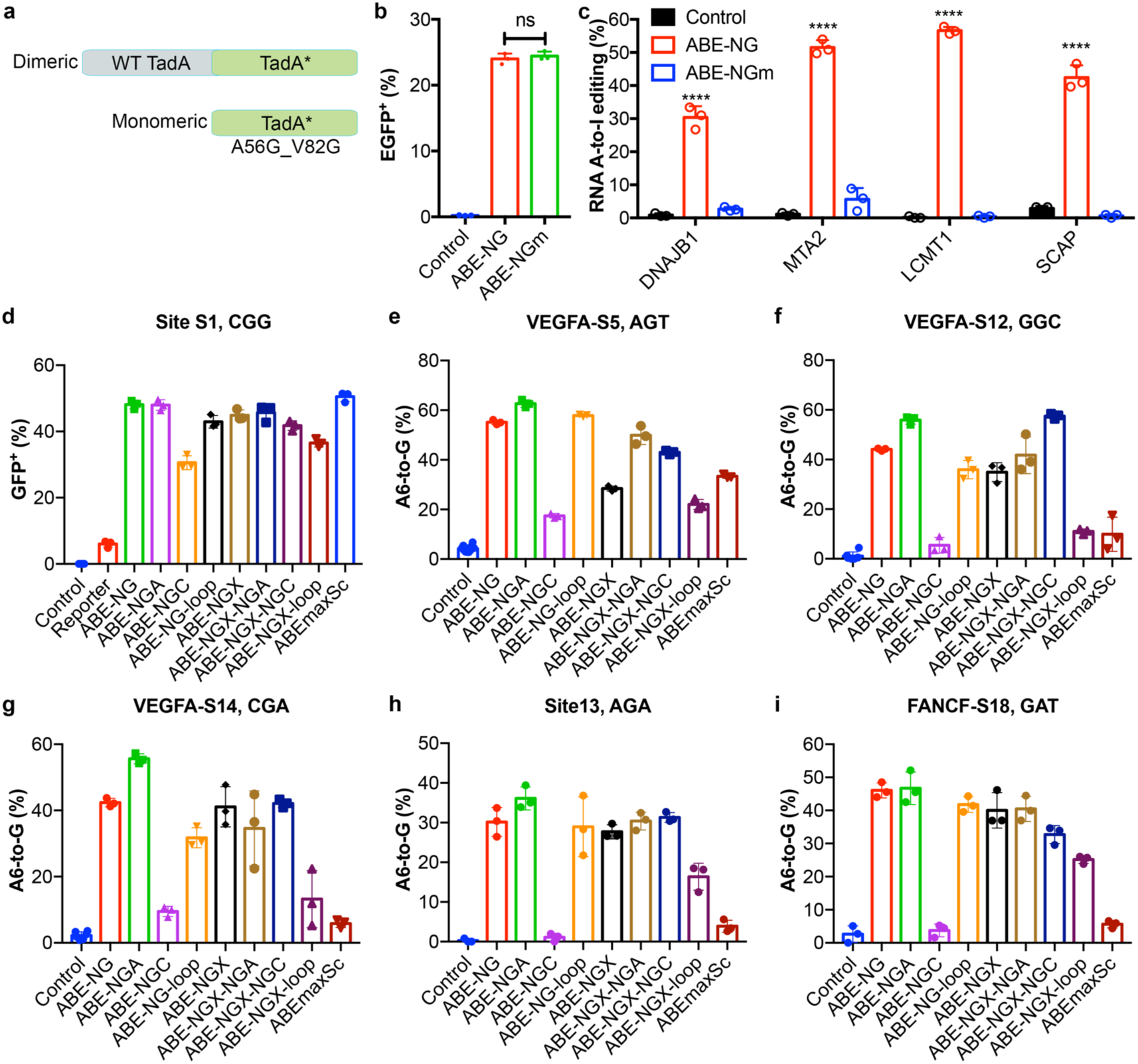
Re-engineering ABE-NG. **a**, Schematics of the adenine deaminase domain used in ABE-NG and ABE-NGm. **b**, ABE-NG and ABE-NGm (the monomeric TadA* with A56G and V82G mutations fused with SpCas9-NG nickase) showed comparable on-target DNA editing activity as assayed by the reporter assay. ns, not statistically significant (one-way ANOVA). **c**, Quantification of the off-target RNA editing (A-to-I) activities on four RNA adenines previously identified as being efficiently modified by ABEmax. ****p < 0.0001 (one-way ANOVA). **d**, Comparison of the base editing efficiency of different ABE variants on site S1 with CGG PAM using the fluorescent reporter assay. **e-i**, Determination of the base editing efficiencies of different ABE variants on five different target sites with NGH or GAT PAM.

Second, we further engineered the PAM-interacting domain of ABE-NG to improve the targeting efficiency at sites with a 5’-NG PAM. We hypothesize that the targeting property of ABE-NG can be modified by combining the mutations in SpCas9-NG (R1335V/L1111R/D1135V/G1218R/E1219F/A1322R/T1337R) with other mutations rationally designed to target different PAM sequences such as those in xCas9(3.7) (A262T/R324L/S409I/E480K/E543D/M694I/E1219V), VQR (D1135V/R1335Q/T1337R), VRER (D1135V/G1218R/R1335E/T1337R) and the loop sequence in ScCas9 (amino acids 367-376). We generated seven new ABE variants with different combinations of the aforementioned variants (details are provided in **Supplementary Table S1**) and compared their base editing activities at six different loci with those of ABE-NG and ABEmaxSC. While all variants except ABE-NGC (containing all NG mutations plus R1335E) performed similarly at the NGG site (**Fig. 3d**), we observed that ABE-NGA (carrying all NG mutations plus R1335Q) had higher editing efficiency than ABE-NG at NGH sites (**Fig. 3e-h**). Interestingly, ABE-NGA and ABE-NGX-NGC (carrying the xCas9(3.7) mutations, NG mutations and R1335E) worked equally well at the NGC site. The ABE-NG and ABE-NGA also edited the site containing a 5’-GAT PAM with high efficiency (**Fig. 3i**). Since ABE-NGA is generally superior to ABE-NG and other variants at NGH sites, we chose ABE-NGA for further *in vivo* studies, and designated the final improved ABE-NGA as iABE-NG, which contains the monomeric TadA*_A56G_V82G.

Finally, we replaced the promoter meCMV with a muscle-specific promoter (truncated MHCK7) to drive the expression of iABE-NG. Interestingly, we had lower AAV9 production yield for our original AAV9-ABE-NG construct than other AAV9 preparations produced at the same time. Although requiring further vigorous examination, we suspected that the ABE-NG expression in the packaging cells during vector production may have a negative impact on the yield. Indeed, we consistently obtained good yield for our new vector AAV9-iABE-NG (designed as AAV9-iNG).

### Systematic delivery of an improved ABE-NG leads to widespread cardiac dystrophin restoration

We treated a cohort of *mdx*^*4cv*^ mice with AAV9-iNG (1×10^14^ vg/kg) through a single tail vein injection at 5 weeks of age. A subset of the mice was sacrificed at 5 weeks after AAV9-iNG administration. Dystrophin was found to be widely rescued in *mdx*^*4cv*^ heart (**Fig. 4a** and **Supplementary Figs. S5-11**). Quantification of the entire heart sections showed that 41.9 ± 10.5% cardiomyocytes of *mdx*^*4cv*^ mice became dystrophin positive following systematic AAV9-iNG treatment (N=5) while the control *mdx*^*4cv*^ hearts were essentially dystrophin negative (0.03 ± 0.02%, N=4) (**Fig. 4b**). Dystrophin was also rescued in skeletal muscles (diaphragm and gastrocnemius) of *mdx*^*4cv*^ mice treated with AAV9-iNG, however, the recovery was less efficient as compared to that in the heart (**Fig. 4c, d**, and **Supplementary Fig. S12**). Western blot analysis showed similar results (**Fig. 4e, f**, and **Supplementary Fig. S13**). The editing efficiency of the dystrophin gene was estimated as described above. The AAV9-iNG treated *mdx*^*4cv*^ hearts showed an average 32.6 ± 2.0% T-to-C editing at the premature stop codon (**Fig. 4g, h**).

**Fig. 4:**
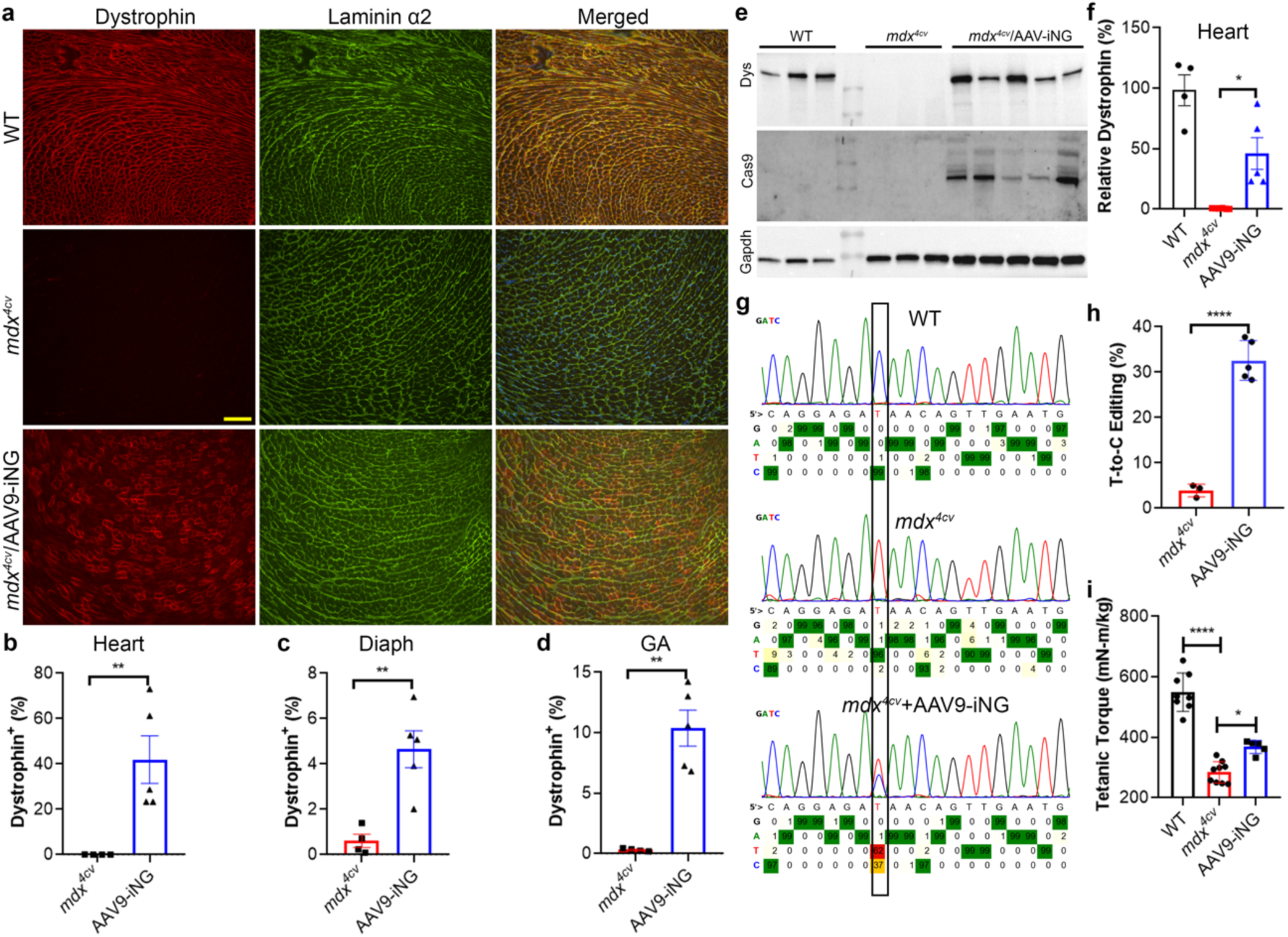
Short-term systemic administration of AAV9-iNG in *mdx*^*4cv*^ mice. **a**, Dystrophin and laminin-α2 co-immunostaining of heart sections from WT and *mdx*^*4cv*^ mice (10 weeks of age) with or without tail vein injection of AAV9-iNG (1×10^14^ vg/kg). Scale bar: 100 µm. **b-d**, Quantification of dystrophin-positive fibers in the heart (**b**), diaphragm (Diaph, **c**) and gastrocnemius (GA, **d**) muscles. **e**, Western blot analysis of heart homogenates with anti-dystrophin, Cas9 and Gapdh antibodies. The WT muscle lysates were loaded at 5 µg/lane while the *mdx*^*4cv*^ muscle lysates were loaded at 25 µg/lane. **f**, Densitometry quantification of Western blot data for the heart muscles. *p < 0.05 (one-way ANOVA). **g**, Representative sequencing trace of dystrophin transcripts of WT and *mdx*^*4cv*^ mouse hearts with or without AAV9-iNG treatment. **h**, Quantification of the targeted T-to-C editing efficiency in the *mdx*^*4cv*^ mouse hearts as assayed by sequencing of dystrophin transcripts. ****p < 0.0001. **i**, Tetanic torque measurements of the posterior compartment muscles. *p < 0.05 (one-way ANOVA).

To test if systemic AAV9-iNG delivery could improve muscle contractility, the mice were examined for muscle force production at 8 weeks of age as described above. While the untreated *mdx*^*4cv*^ mice produced significantly reduced torque as compared to the WT controls, systemic delivery of AAV9-iNG significantly improved the tetanic torque in *mdx*^*4cv*^ mice (**Fig. 4i**).

### Systematic delivery of AAV9-iNG leads to long-term, near complete dystrophin rescue in dystrophic hearts

A group of *mdx*^*4cv*^ mice treated with intravenous administration of AAV9-iNG at 5 weeks of age were kept for over 10 months to study the long-term impact of systemic ABE editing therapy. Surprisingly, a near complete dystrophin restoration was observed in the hearts of all four treated *mdx*^*4cv*^ mice (**Fig. 5a, b** and **Supplementary Figs. S14-19**). Western blot analysis showed similar results (**Fig. 5c, d**). Sequencing of dystrophin mRNA revealed 84.6 ± 2.6%T-to-C editing at the targeted mutation site (**Fig. 5e, f**). The editing efficiencies in the skeletal muscles were substantially lower than those in the hearts (**Supplementary Fig. S20**).

**Fig. 5:**
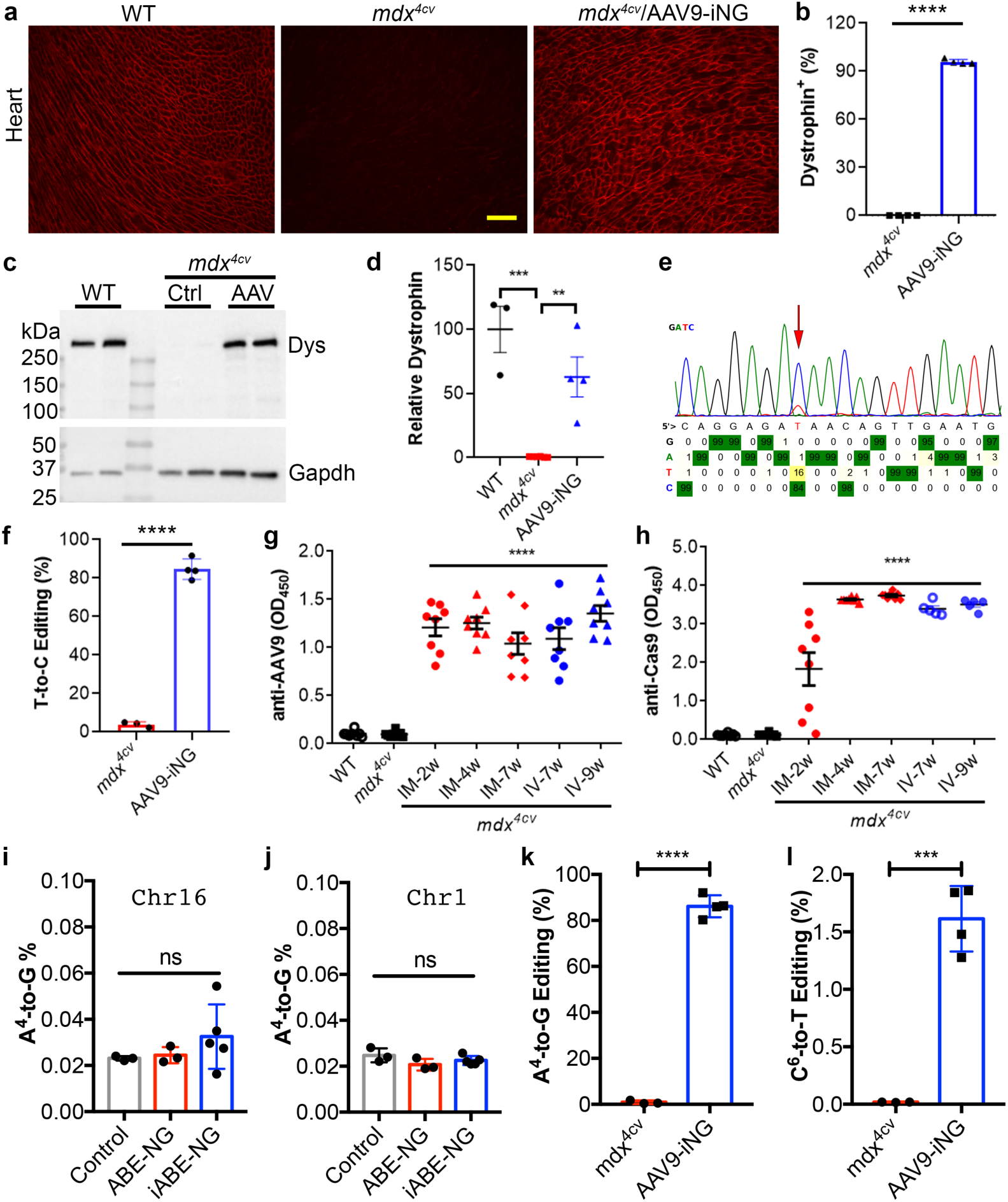
Long-term efficacy, host response and toxicity studies of AAV9-iNG therapy in *mdx*^*4cv*^ mice. **a**, Dystrophin immunostaining of heart sections from WT and *mdx*^*4cv*^ mice (10 months of age) with or without tail vein injection of AAV9-iNG (1×10^14^ vg/kg) at 5 weeks old. Scale bar: 100 µm. **b**, Quantification of dystrophin-positive fibers in the heart of *mdx*^*4cv*^ mice. ****p < 0.0001 (*t*-test) **c**, Western blot analysis of heart homogenates with anti-dystrophin and Gapdh antibodies. **d**, Densitometry quantification of Western blot data for the hearts. ****p < 0.0001; **p < 0.01 (one-way ANOVA) **e**, Representative sequencing trace of dystrophin transcripts of *mdx*^*4cv*^ mouse heart receiving AAV9-iNG treatment. The targeted T-to-C conversion is marked by a red arrow. **f**, Quantification of the targeted T-to-C editing efficiency in the *mdx*^*4cv*^ mouse hearts as assayed by sequencing of dystrophin transcripts. ****p < 0.0001 (*t*-test). **g-h**, Host immune response to AAV9 and ABE-NG or iNG (anti-Cas9). IM, intramuscular injection; IV, intravenous injection. ****p < 0.0001 (one-way ANOVA) compared to WT or *mdx*^*4cv*^. **i-j**, Quantification of deep sequencing reads of the genomic DNA PCR amplicons of the chromosome 16 off-target site (**i**) or the chromosome 1 off-target site (**j**) from Neuro-2a cells transfected with ABE-NG, iABE-NG or control plus the gRNA. ns, not significant (one-way ANOVA). **k**, Quantification of the A^4^-to-G editing in *mdx*^*4cv*^ mice treated with or without AAV9-iNG. **** p < 0.0001 (student’s *t*-test). **l**, Quantification of the bystander C^6^-to-T editing in *mdx*^*4cv*^ mice treated with or without AAV9-iNG. *** p < 0.001 (student’s *t*-test).

Previous studies from others and us showed that AAV-mediated delivery of CRISPR/Cas9 into neonatal mice resulted in humoral immune responses to AAV capsid but not Cas9^14, 57^. In contrast, AAV-mediated delivery of CRISPR/Cas9 into adult mice evoked robust anti-Cas9 immunity^14^. Serum samples were collected to analyze the host immune responses to the AAV9 capsid and the base editor ABE-NG. Intramuscular injection of AAV9-NG into 5-6 weeks old *mdx*^*4cv*^ mice produced robust anti-AAV9 capsid (**Fig. 5g**) and anti-Cas9 antibodies (**Fig. 5h**) at 2 weeks after injection. The anti-AAV9 titers were similar at different time points from 2 to 7 weeks post intramuscular injection and from 7 to 9 weeks post intravenous injection (**Fig. 5g**). The anti-Cas9 antibody titers showed a large variation among mice at 2 weeks after intramuscular injection, but all increased to peak by 4 weeks (**Fig. 5h**).

We further examined the liver toxicity by measuring serum aspartate aminotransferase (AST) and alanine aminotransferase (ALT), and kidney toxicity by measuring blood urine nitrogen (BUN). As compared to WT mice, the *mdx*^*4cv*^ mice showed elevated AST and ALT (**Supplementary Fig. S21a, b**). However, treatment of *mdx*^*4cv*^ mice with AAV9-iNG did not further increase the serum levels of AST and ALT (**Supplementary Fig. S21a, b**), at either 8 weeks or 10 months of age. Measurement of BUN did not find significant changes in the treated or untreated *mdx*^*4cv*^ mice (**Supplementary Fig. S21c**).

### AAV9-iNG treatment exhibits low off-target activities *in vivo* and *in vitro*

One concern with ABE-mediated gene correction is the potential off-target activities such as gRNA mismatch tolerance, bystander editing, and off-target RNA editing. Previous studies showed that ABE can tolerate 1-2 mismatches between the gRNA and its target sites ^58^. Prediction by Cas-OFFinder showed that one site on chromosome 16 (Chr16_OT) has only one mismatch, two other sites have two mismatches and 55 sites have three mismatches. The Chr16_OT differs from the *mdx*^*4cv*^ target sequence by only one C at position 12 (**Supplementary Fig. S22a**). We transfected Neuro-2a cells with ABE-NG or iABE-NG plus the gRNA, amplified the Chr16_OT by PCR and subjected the amplicon to next generation sequencing (NGS). As shown in **Fig. 5i**, we did not observe significant editing of the A4 in either ABE-NG or iABE-NG transfected cells. Similarly, we analyzed the off-target site on chromosome 1 (Chr1_OT), which differs from the *mdx*^*4cv*^ target sequence by an A at position 2 and a G at position 20 (**Supplementary Fig. S22a**). Again, we found that ABE-NG or iABE-NG did not edit the A4 at Chr1_OT (**Fig. 5j**). These data suggest that ABE-NG and iABE-NG show low tolerance to mismatches.

Next, we analyzed the potential bystander editing at the on-target *mdx*^*4cv*^ locus in the mice treated with AAV9-iNG. Since the 10-month treated mouse hearts showed a high level of dystrophin rescue, we first determined the on-target editing efficiency in these mouse hearts by NGS. As mouse hearts contain multiple different cell types, we predicted that analysis of the genomic DNA PCR products would significantly underestimate the editing efficiency. To verify this, we performed NGS of the genomic DNA PCR products from two mouse hearts receiving AAV9-iNG and exhibiting high dystrophin rescue, and detected an up to 11% edits at A4. Thus, we sequenced the RT-PCR products to estimate the editing efficiency at the on-target *mdx*^*4cv*^ locus. The A at position 4 (corresponding to the T within the premature stop codon in the coding strand) was converted to G with high efficiency from all four mouse hearts (**Supplementary Fig. S22b**). On average, 86.2 ± 2.4 % A-to-G conversion was measured (**Fig. 5k**). At the *mdx*^*4cv*^ target site, there was only one A within the editing window of 4-8, disallowing us to analyze the bystander A-to-G editing at this site. Another type of undesired ABE-mediated genome edits at an on-target locus is ABE-dependent cytosine-to-uracil conversion resulting in C•G to T•A mutation at that site ^55, 59, 60^. We found that C6 at the *mdx*^*4cv*^ target site was edited above background with an average efficiency of 1.6 ± 0.1 % (**Fig. 5l**).

## Discussion

Collectively, our study has shown that AAV9-mediated delivery of a rationally improved NG-targeting ABE allowed systemic restoration of dystrophin expression and functional improvement. The editing efficiency in the heart was extraordinarily high in *mdx*^*4cv*^ mice following systemic delivery of AAV9-iNG and over 90% of cardiomyocytes were corrected to express dystrophin in *mdx*^*4cv*^ hearts at 10 months of age after a single intravenous administration of AAV9-iNG at 5 weeks old. There was no obvious toxicity detected following AAV9-iNG treatment, despite the host immune response to the AAV9 capsid and ABE. This has tremendous implication for base correction of genetic cardiomyopathies.

The iABE-NG could be broadly applied to correct DMD mutations and potentially many other disease-causing mutations. Analysis of the ClinVar database showed that over 100 of the 174 total G>A or T>C point mutations for DMD could be targeted for repair by at least one of the ABEs (iABE-NG). The recent advances in engineering Cas9 variants with non-G PAM^33, 34^ would further increase our targeting capacity. Moreover, the ABE editing could be designed to induce skipping of mutant exons via targeting the canonical splicing donor or acceptor ^61, 62^, thus further broadening the applicability of ABE editing therapy for a larger population of DMD.

It is of interest that the mice at ten months after AAV9-iNG delivery appeared to show significantly higher dystrophin rescue than the mice at 10 weeks after the treatment. One plausible explanation is that the DMD cardiomyocytes with restored dystrophin expression may gain advantage for selective survival and regeneration during the development stages after delivery of AAV9-iNG. Additionally, transduced cardiomyocyte-derived extracellular vesicles may deliver genetic materials such as transcripts encoding iABE-NG into proximal un-transduced cardiomyocytes and confer base editing in those cells.

Our study has also shown that systemic delivery of AAV9-iNG resulted in dystrophin restoration in skeletal muscles and functional improvement. As compared to cardiomyocytes, the editing efficiency in skeletal muscles were substantially lower. An obvious explanation is that AAV9 has higher tropism towards cardiomyocytes than skeletal muscles^63^. However, other mechanisms may also be responsible for the lower editing efficiency in skeletal muscles. For example, the dystrophic and inflammatory microenvironment in skeletal muscles may pose further strains on AAV9 delivery and base editing. In addition, targeting muscle satellite cells may be required to improve the overall editing outcomes in skeletal muscle as they are constantly activated to replace injured skeletal muscle in DMD. Although AAV9 has been shown to transduce muscle satellite cells, the efficiency is relatively low^64-67^. Moreover, the use of a muscle-specific promoter would probably further reduce the base editing in muscle satellite cells in our present study. Future studies are required to improve the base editing efficiency in dystrophic skeletal muscles.

Recently, Liu and colleagues employed a similar intein PTS approach to split ABE and CBE at the Cys 574 split site of Cas9 and packaged into AAV for *in vivo* delivery, and showed ∼ 20% editing efficiency in heart^68^, significantly lower than the editing efficiency achieved in our study. Several differences between these studies may explain the different editing efficiencies observed. First, the intein used in our study (Gp41-1) has a superfast kinetics as compared to that used by Levy *et al*.., which may speed the assembly of full-length ABE. Second, each half of the AAVs carries a gRNA-expressing cassette while in the study by Levy et al., gRNA is only present in one of the AAV constructs. Previous studies have shown that the gRNA dosage plays an important role in the editing efficiency. Third, we used a single mutant TadA* domain, which was made to eliminate the off-target RNA editing activity, while Levy et al., used the original dimeric ABE7.10 in their study, which was shown to have high off-target RNA editing activity. Fourth, the promoters used in these studies were also different, which may drive different expression levels of ABE in heart tissues. Finally, the intrinsic difference in the gRNAs and ABE variants may have impacts on the overall editing outcomes. Nevertheless, the exceptionally high editing efficiency achieved in adult dystrophic mice suggests that our optimized AAV-iNG vectors could be used for future clinical applications.

## Data availability

The authors declare that all data supporting the findings of this study are available in the article and its Supplementary Information files or are available from the corresponding author on reasonable request.

## Acknowledgements

The authors thank the Viral Vector Core at the Nationwide Children’s hospital for producing the AAV, the Analytical Cytometry Shared Resource of the Ohio State University Comprehensive Cancer Center for FACS, the Genomics Shared Resource of the Ohio State University Comprehensive Cancer Center for sequencing, and Katiri Snyder and Natalie Sell for assisting in tail vein injection. R.H. is supported by US National Institutes of Health grant (R01 HL116546) and a Parent Project Muscular Dystrophy award.

## Author information

These authors contributed equally: Li Xu, Chen Zhang.

## Contributions

R.H. conceived the study and wrote the manuscript. L.X., C.Z. and P.W. performed the molecular cloning, cell culture studies, immunofluorescence and Western blot experiments and analyzed the data. Y.G. and C.Z. performed the ELSA experiments, assisted the AAV injections and collected blood samples. H.L. and C.Z. maintained the mouse colony and coordinated mouse tissue collections. P.W. and L.X. performed FACS experiments and analyses. W.D.A., C.Z. and R.H. performed *in vivo* muscle contractility experiments. R.H. wrote the manuscript.

