## Supplemental Data for "Efficient precise *in vivo* base editing in adult dystrophic mice"

**First author's surname**: Xu

**Short title**: *in vivo* base editing of DMD mice

Li Xu[1†], Chen Zhang[1†], Haiwen Li[1], Peipei Wang[1], Yandi Gao[1], Peter J. Mohler[2], Nahush A. Mokadam[1], Jianjie Ma[1], William. D. Arnold[3], Renzhi. Han[1*]

### Supplemental Material

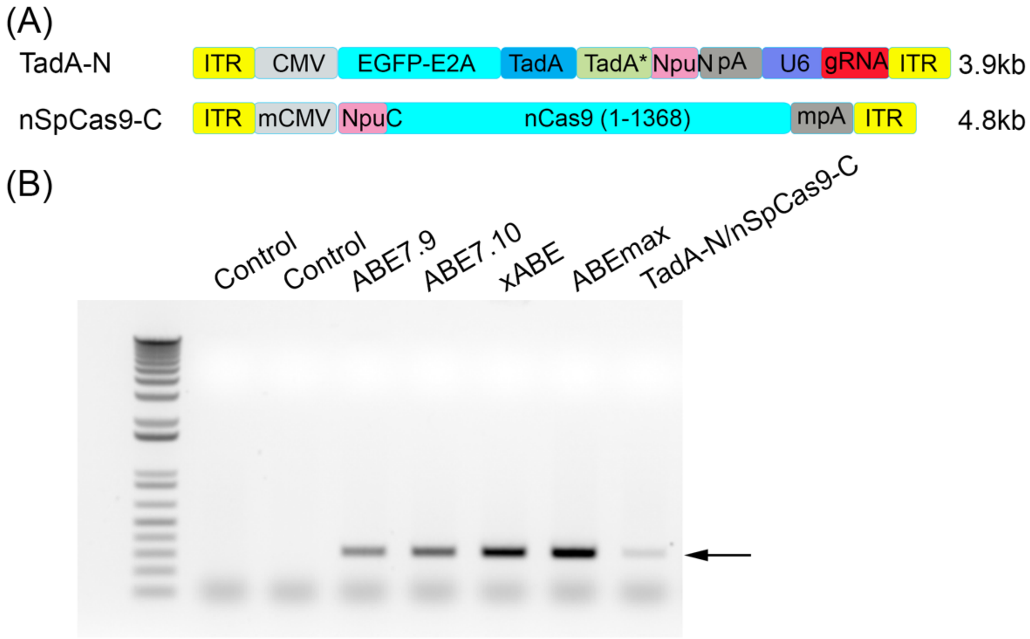


**Fig. S1**. The intein split of ABEmax had relatively low editing activity. (A) Schematics showing the two halves of intein-split ABEmax. The TadA-TadA* was fused with Npu intein N-terminal fragment and SpCas9 nickase (nSpCas9) was fused with Npu intein C-terminal fragment. (B) Genomic DNA PCR analysis of HEK293 cells at 5 days after transfection with S2-gRNA and different versions of ABEs.

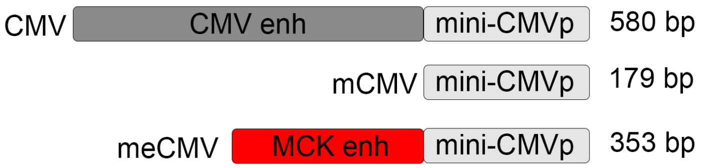


**Fig. S2**. Schematics of meCMV promoter. The meCMV promoter is composed of a 179-bp mini-CMV core promoter and 174-bp mouse creatine kinase (MCK) enhancer sequence.

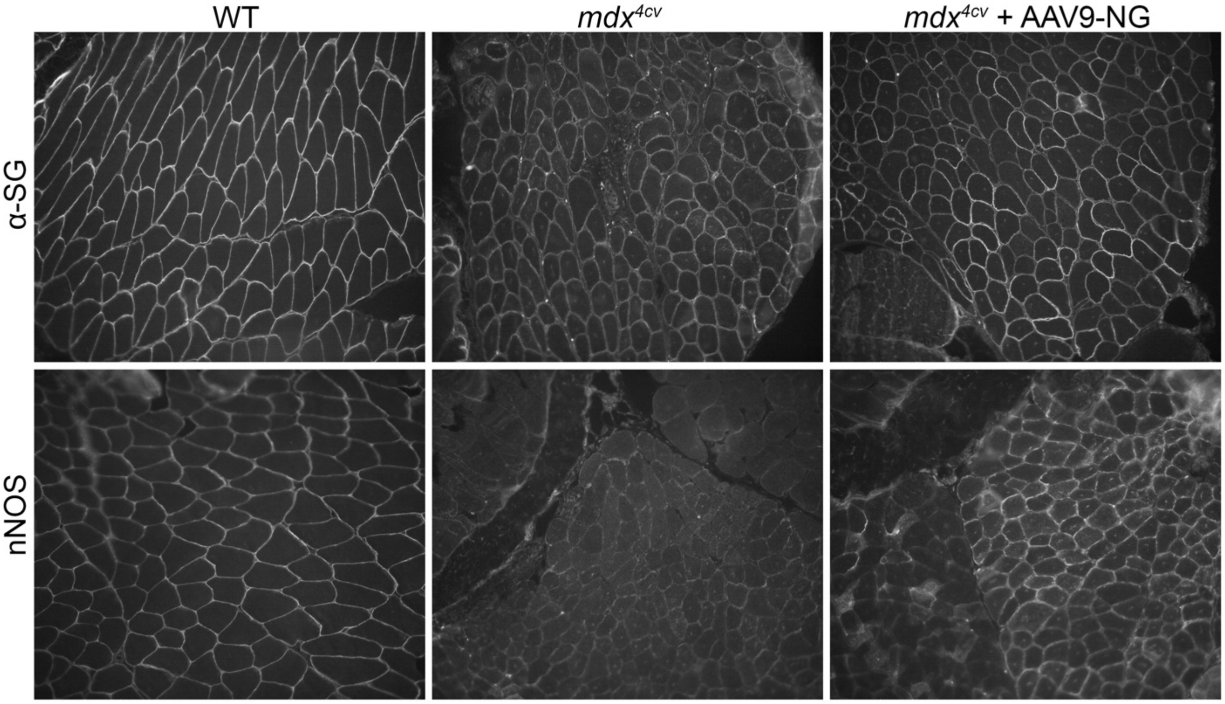


**Fig. S3**. Immunofluorescence staining of gastrocnemius muscle sections with anti-α-sarcoglycan (α-SG) and anti-neuronal nitric oxide synthase (nNOS). Scale bar: 200 μm.

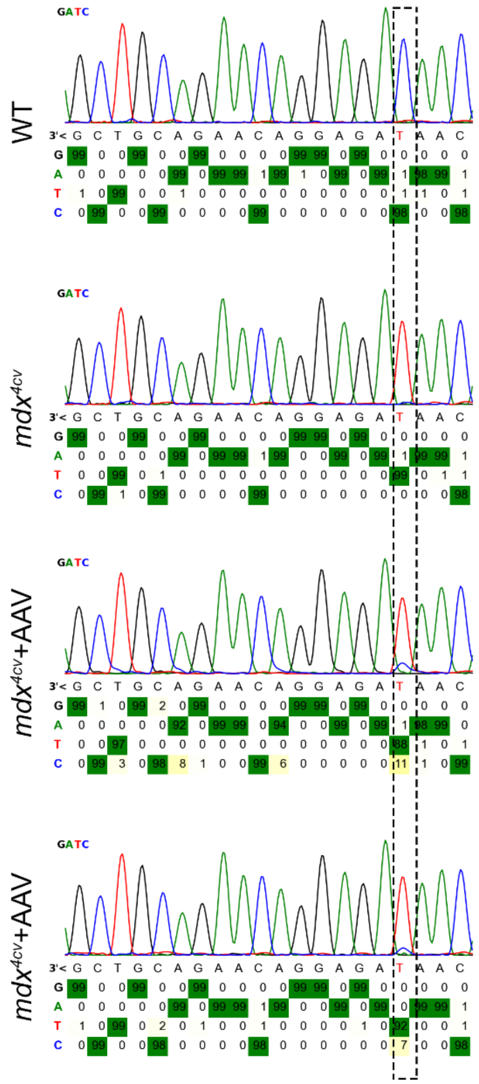


**Fig. S4**. Sanger sequencing traces of dystrophin cDNA showing the T-to-C editing in *mdx*$^{4cv}$ mice receiving intramuscular injection of AAV9-NG.

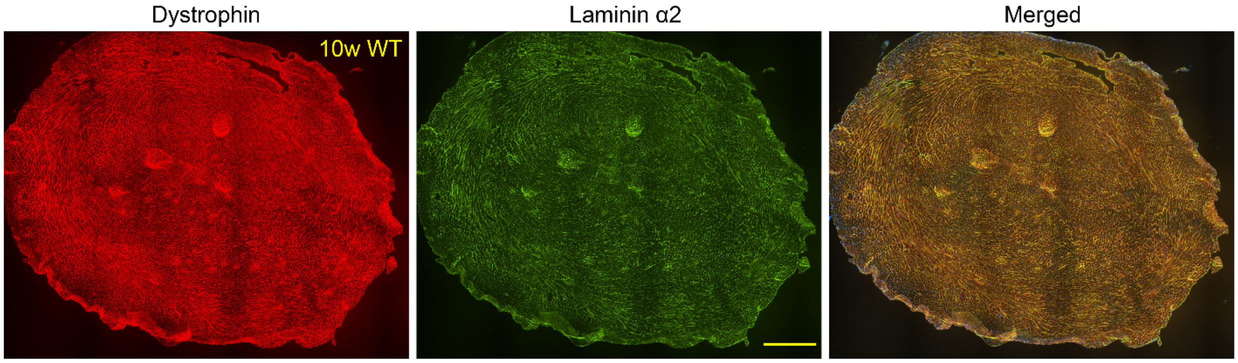


**Fig. S5.** Stitched large images showing dystrophin and laminin-α2 immunostaining of the entire heart sections of a WT mouse at 10 weeks of age. Mouse number is shown in yellow. Scale bars: 0.5 mm.

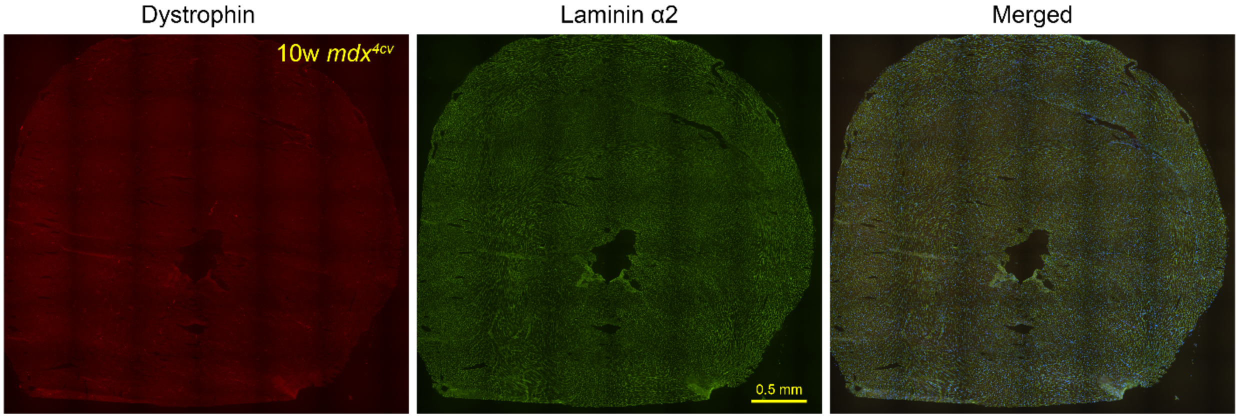


**Fig. S6.** Stitched large images showing dystrophin and laminin-α2 immunostaining of the entire heart sections of a control *mdx*$^{4cv}$ mouse at 10 weeks of age. Mouse number is shown in yellow. Scale bars: 0.5 mm.

**Fig. S7-11.** Stitched large images showing dystrophin and laminin-α2 immunostaining of the entire heart sections of five individual *mdx*$^{4cv}$ mice five weeks after intravenous injection of AAV9-iNG at 5 weeks of age. Mouse number is shown in yellow. Scale bars: 0.5 mm.

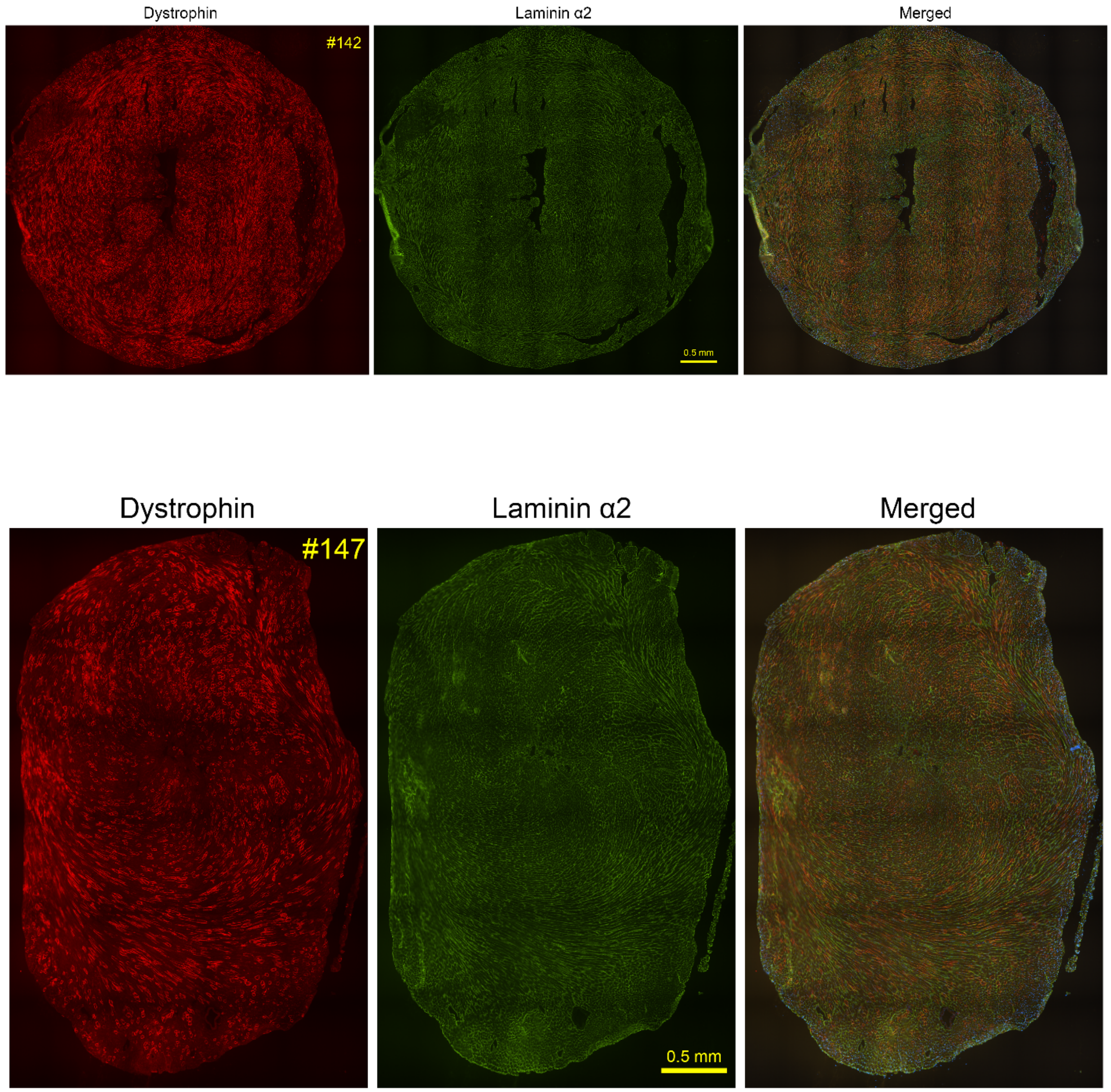

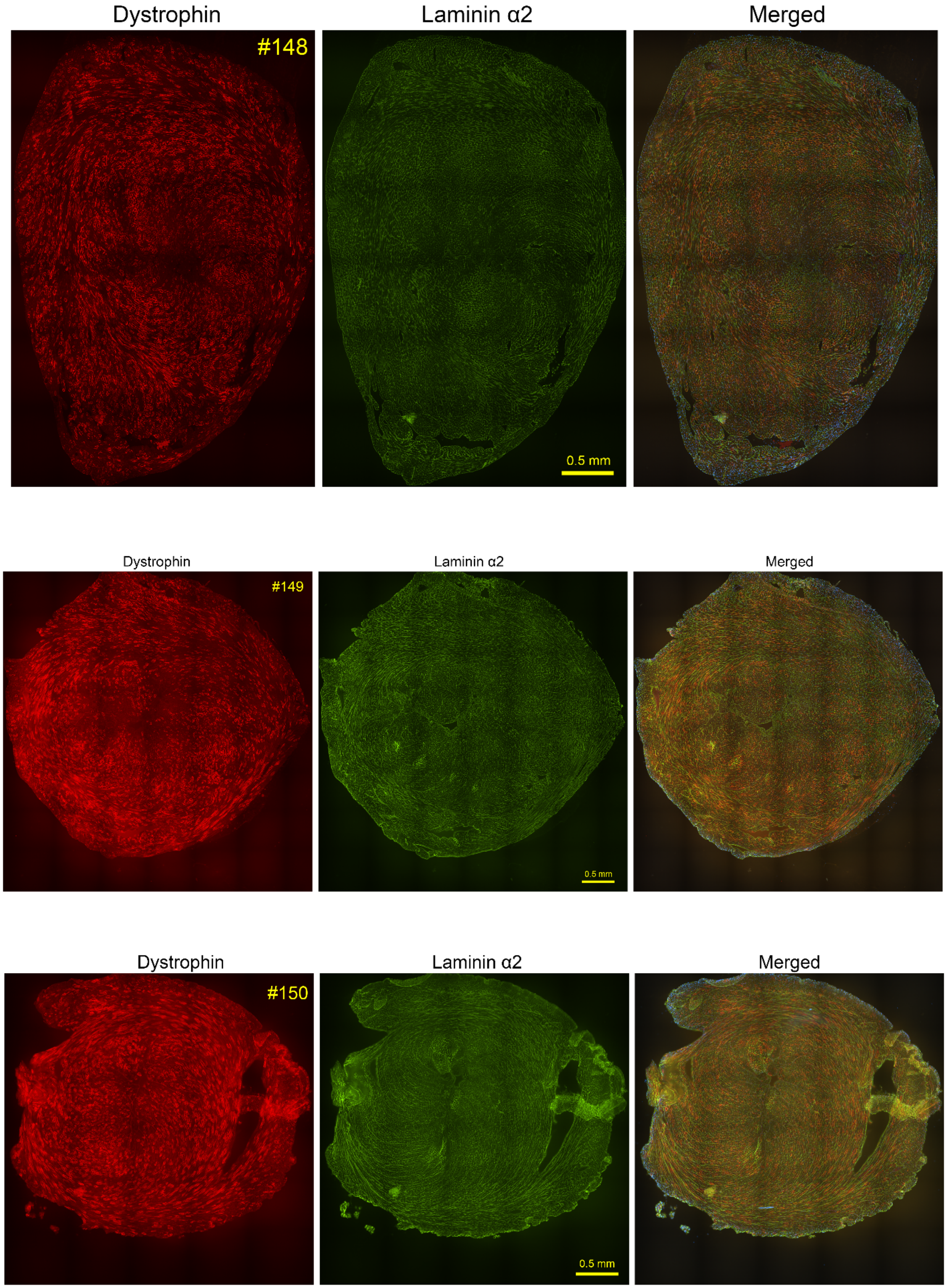
Dystrophin
Laminin α2
Merged
#148
0.5 mm
Dystrophin
Laminin α2
Merged
#149
0.5 mm
Dystrophin
Laminin α2
Merged
#150
0.5 mm

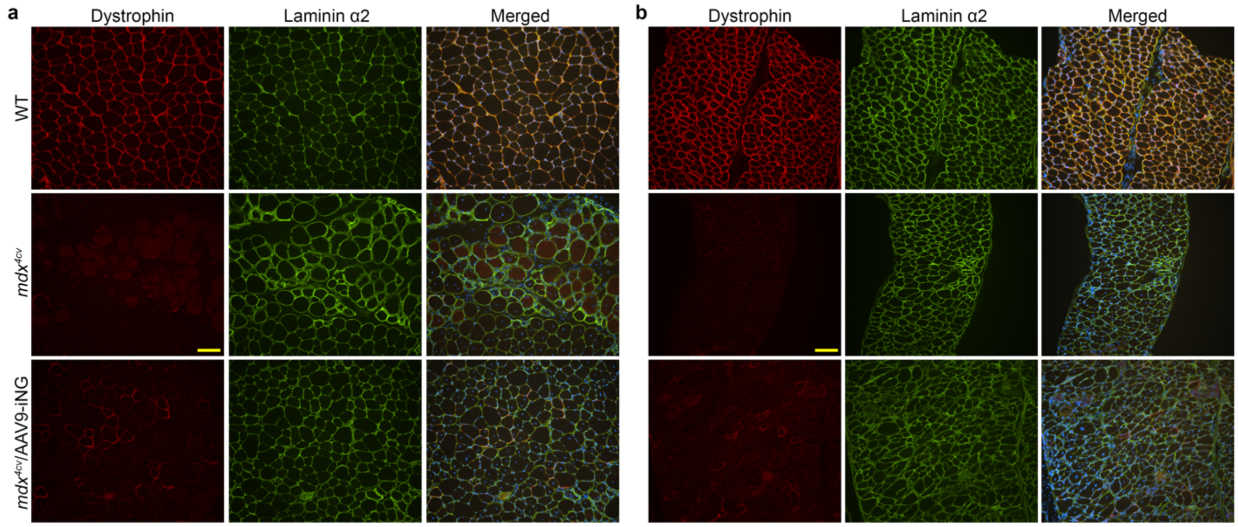


**Fig. S12**. Immunofluorescence staining of dystrophin and laminin α2 in the gastrocnemius (**a**) and diaphragm (**b**) muscles from WT and *mdx*$^{4cv}$ treated with or without tail vein injection of AAV9-iNG. Scale bar: 100 µm.

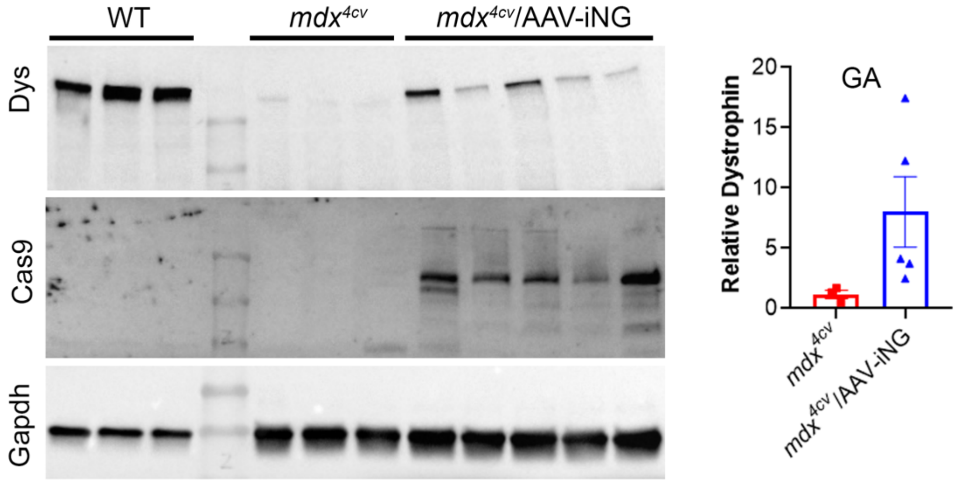


**Fig. S13**. Western blot analysis of gastrocnemius muscles from WT and *mdx*$^{4cv}$ treated with or without tail vein injection of AAV9-iNG.

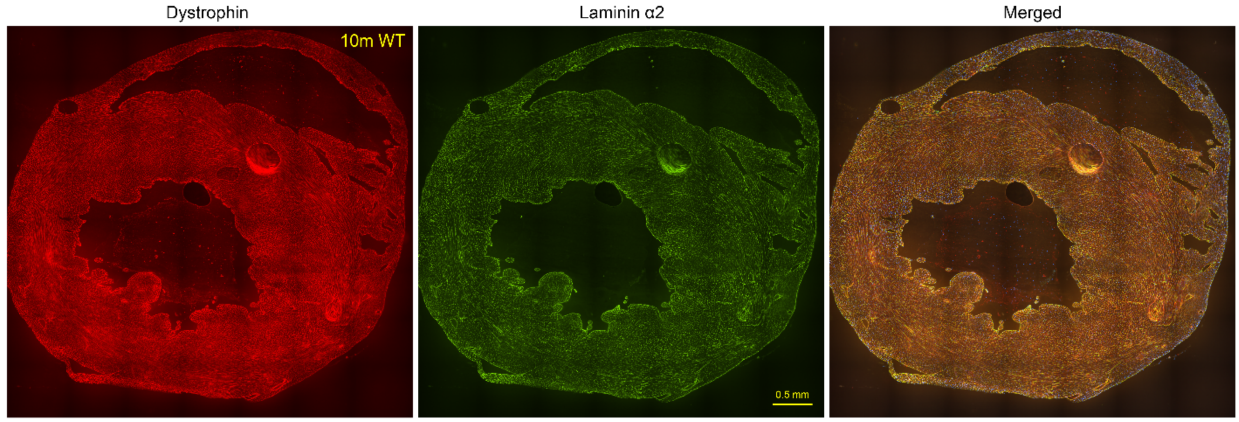


**Fig. S14.** Stitched large images showing dystrophin and laminin-α2 immunostaining of the entire heart sections of a WT mouse at 10 months of age. Mouse number is shown in yellow. Scale bars: 0.5 mm.

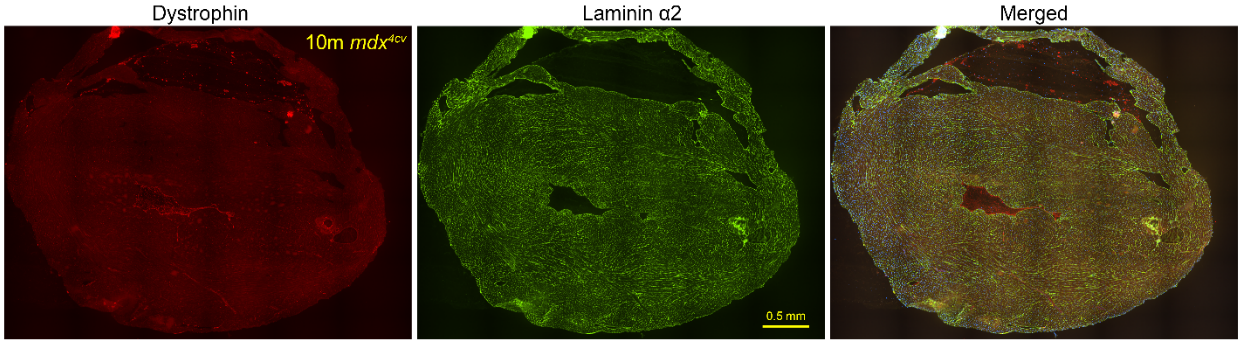


**Fig. S15.** Stitched large images showing dystrophin and laminin-α2 immunostaining of the entire heart sections of a control *mdx*$^{4cv}$ mouse at 10 months of age. Mouse number is shown in yellow. Scale bars: 0.5 mm.

**Fig. S16-19.** Stitched large images showing dystrophin and laminin-α2 immunostaining of the entire heart sections of four individual *mdx*$^{4cv}$ mice 9-10 months after intravenous injection of AAV9-iNG at 5 weeks of age. Mouse number is shown in yellow. Scale bars: 0.5 mm.

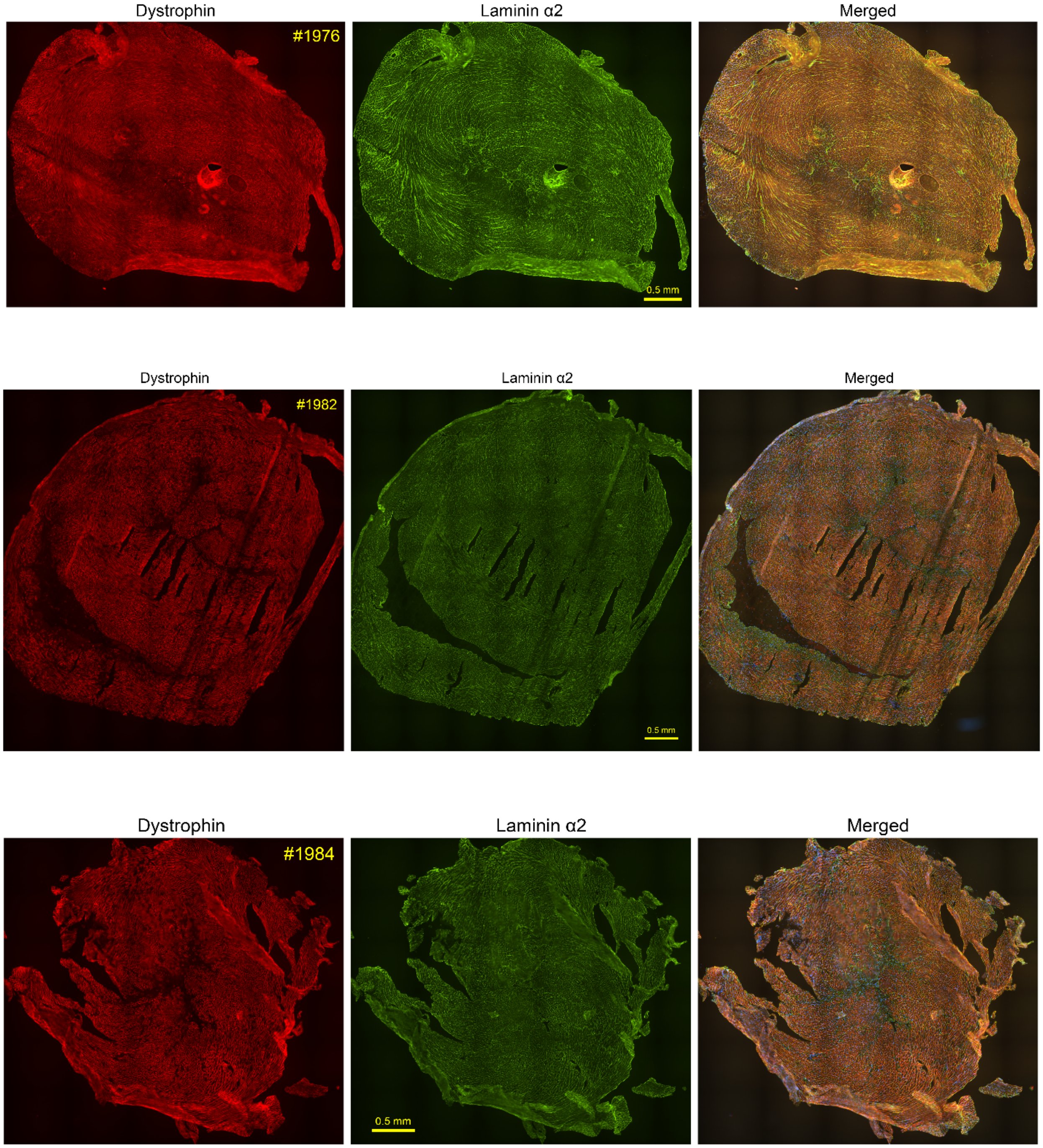

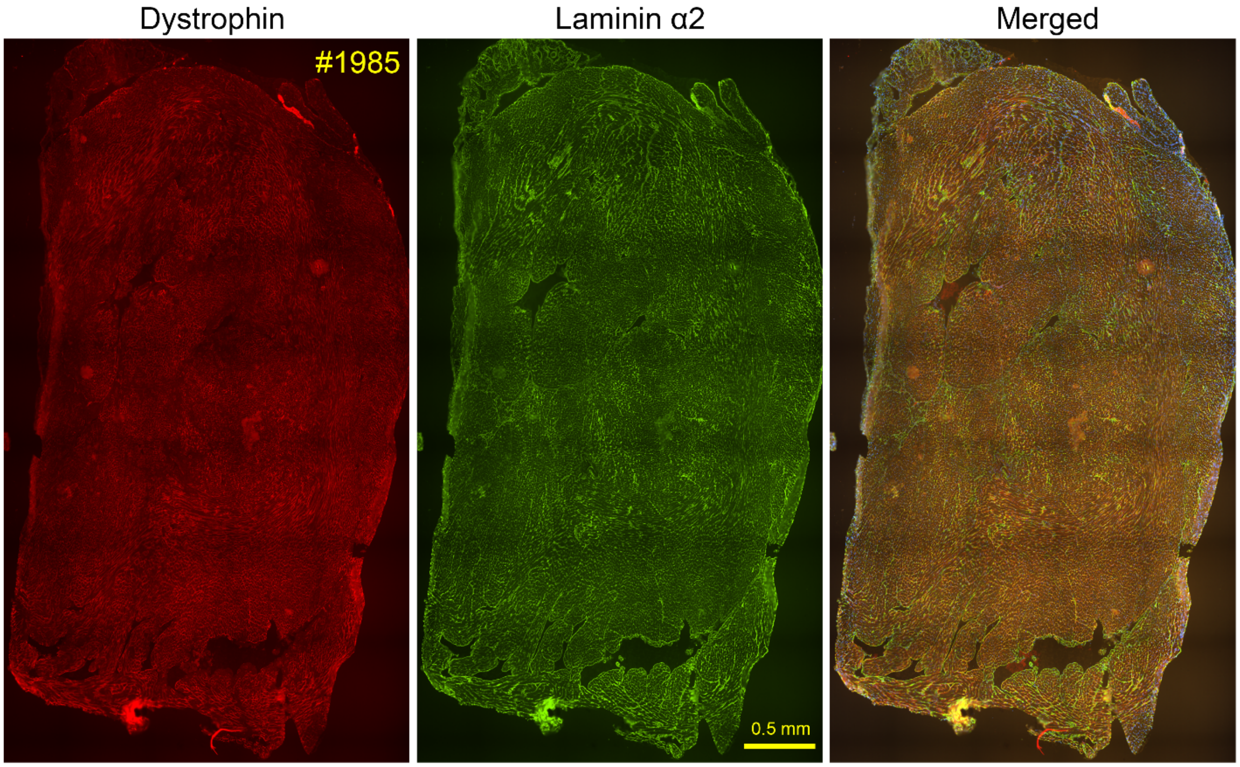

Dystrophin
Laminin α2
Merged
#1985
0.5 mm

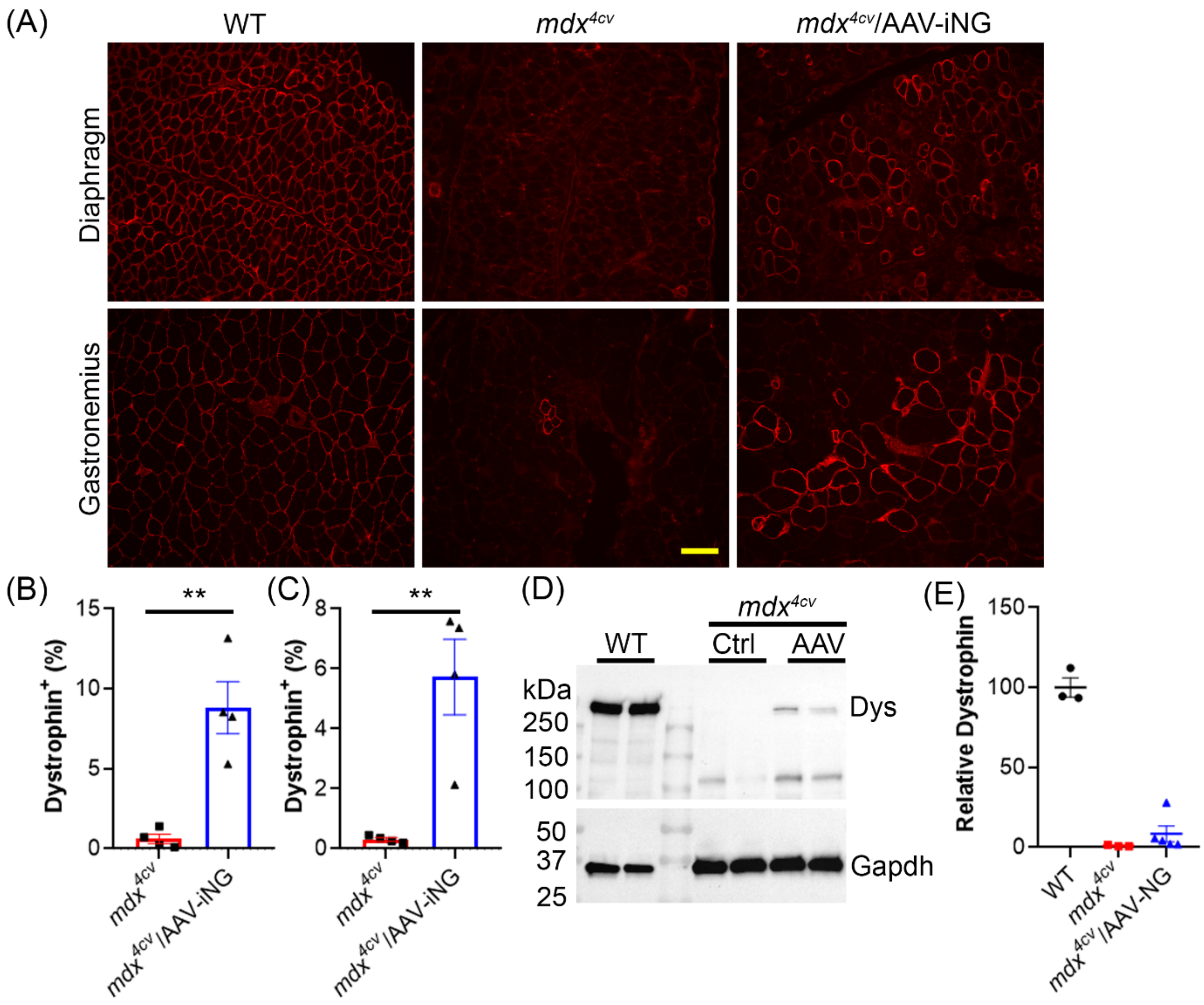


**Fig. S20**. Restoration of dystrophin expression in the skeletal muscles of 10-month-old *mdx*$^{4cv}$ mice after tail vein injection of AAV9-iNG at 5 weeks of age. (A) Immunofluorescence staining of dystrophin in diaphragm and gastrocnemius muscles of WT and *mdx*$^{4cv}$ mice with or without systemic AAV9-iNG delivery. (B-C) Quantification of dystrophin+ fibers in diaphragm (B0 and gastrocnemius (C) muscles. (D) Western blot of dystrophin expression in gastrocnemius muscles. (E) Quantification of Western blot data.

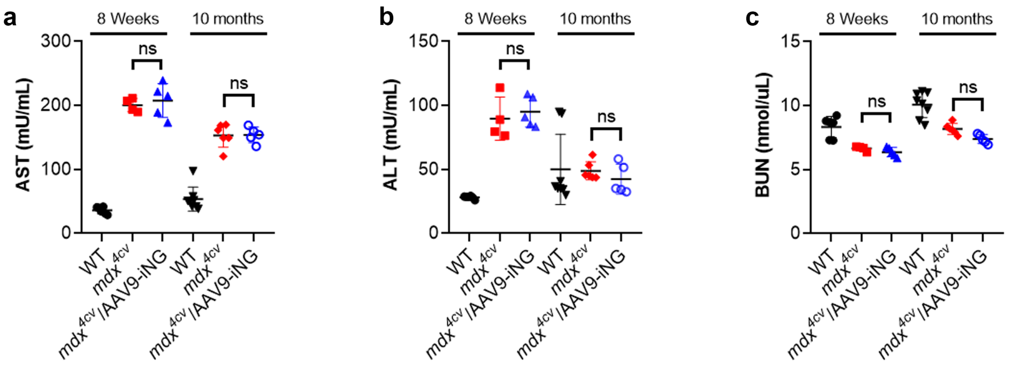


**Fig. S21**. Measurements of serum AST (**a**), ALT (**b**) and BUN (**c**) of mice treated with or without AAV9-iNG. ns, not statistically significant (one-way ANOVA).

**a**

```
 ChrX: GTTATCTCCTGTTCTGCAGC TGT
Chr16: GTTATCTCCTGCTCTGCAGC AGA
 Chr1: GATATCTCCTGTTCTGCAGG AGA
```

**b**

| | | G | T | T | A4 | T | C6 | T | C | C | T | G | T | T | C | T | G | C | A | G | C | T (PAM) | G (PAM) | T (PAM) |
|---|---|---|---|---|---|---|---|---|---|---|---|---|---|---|---|---|---|---|---|---|---|---|---|---|
| Mouse1 | A | 0.05 | 0.01 | 0.00 | 13.64 | 0.01 | 0.03 | 0.03 | 0.05 | 0.02 | 0.02 | 0.02 | 0.01 | 0.01 | 0.00 | 0.03 | 0.08 | 0.01 | 99.90 | 0.02 | 0.01 | 0.01 | 0.03 | 0.01 |
| | C | 0.00 | 0.03 | 0.07 | 0.01 | 0.03 | 97.93 | 0.04 | 99.86 | 99.96 | 0.09 | 0.00 | 0.02 | 0.03 | 99.96 | 0.11 | 0.00 | 99.96 | 0.00 | 0.00 | 99.97 | 0.07 | 0.00 | 0.02 |
| | G | 99.95 | 0.00 | 0.00 | 86.34 | 0.00 | 0.19 | 0.00 | 0.02 | 0.00 | 0.01 | 99.97 | 0.00 | 0.00 | 0.00 | 0.00 | 99.92 | 0.00 | 0.08 | 99.97 | 0.00 | 0.05 | 99.96 | 0.00 |
| | T | 0.00 | 99.96 | 99.92 | 0.02 | 99.95 | 1.84 | 99.93 | 0.06 | 0.02 | 99.89 | 0.00 | 99.97 | 99.97 | 0.03 | 99.86 | 0.00 | 0.03 | 0.01 | 0.02 | 0.02 | 99.87 | 0.01 | 99.97 |
| Mouse2 | A | 0.07 | 0.01 | 0.00 | 14.04 | 0.02 | 0.04 | 0.03 | 0.04 | 0.03 | 0.02 | 0.04 | 0.01 | 0.01 | 0.01 | 0.03 | 0.07 | 0.01 | 99.90 | 0.04 | 0.01 | 0.01 | 0.03 | 0.01 |
| | C | 0.00 | 0.03 | 0.06 | 0.00 | 0.02 | 98.27 | 0.04 | 99.90 | 99.92 | 0.09 | 0.00 | 0.02 | 0.03 | 99.97 | 0.14 | 0.00 | 99.94 | 0.01 | 0.00 | 99.97 | 0.09 | 0.00 | 0.02 |
| | G | 99.92 | 0.00 | 0.00 | 85.93 | 0.00 | 0.21 | 0.00 | 0.00 | 0.03 | 0.01 | 99.96 | 0.00 | 0.00 | 0.00 | 0.01 | 99.93 | 0.00 | 0.08 | 99.94 | 0.00 | 0.01 | 99.96 | 0.00 |
| | T | 0.00 | 99.95 | 99.94 | 0.02 | 99.96 | 1.48 | 99.92 | 0.05 | 0.02 | 99.88 | 0.00 | 99.97 | 99.96 | 0.02 | 99.82 | 0.00 | 0.04 | 0.02 | 0.02 | 0.02 | 99.89 | 0.01 | 99.97 |
| Mouse3 | A | 0.08 | 0.01 | 0.01 | 7.94 | 0.02 | 0.04 | 0.03 | 0.02 | 0.04 | 0.02 | 0.03 | 0.02 | 0.01 | 0.01 | 0.04 | 0.07 | 0.00 | 99.88 | 0.02 | 0.01 | 0.01 | 0.03 | 0.01 |
| | C | 0.00 | 0.06 | 0.06 | 0.01 | 0.05 | 97.74 | 0.07 | 99.97 | 99.94 | 0.09 | 0.00 | 0.02 | 0.03 | 99.98 | 0.15 | 0.00 | 99.95 | 0.01 | 0.00 | 99.96 | 0.09 | 0.00 | 0.02 |
| | G | 99.91 | 0.00 | 0.00 | 92.03 | 0.00 | 0.36 | 0.01 | 0.00 | 0.01 | 0.01 | 99.97 | 0.00 | 0.00 | 0.00 | 0.01 | 99.92 | 0.00 | 0.10 | 99.97 | 0.00 | 0.01 | 99.96 | 0.01 |
| | T | 0.00 | 99.93 | 99.93 | 0.02 | 99.93 | 1.86 | 99.89 | 0.01 | 0.02 | 99.89 | 0.00 | 99.96 | 99.95 | 0.02 | 99.81 | 0.00 | 0.05 | 0.01 | 0.01 | 0.02 | 99.89 | 0.01 | 99.96 |
| Mouse4 | A | 0.10 | 0.01 | 0.00 | 19.63 | 0.02 | 0.03 | 0.04 | 0.03 | 0.04 | 0.02 | 0.05 | 0.01 | 0.01 | 0.01 | 0.03 | 0.08 | 0.01 | 99.87 | 0.03 | 0.01 | 0.01 | 0.04 | 0.01 |
| | C | 0.00 | 0.04 | 0.06 | 0.01 | 0.02 | 98.51 | 0.06 | 99.93 | 99.94 | 0.10 | 0.00 | 0.03 | 0.04 | 99.97 | 0.17 | 0.00 | 99.93 | 0.01 | 0.00 | 99.96 | 0.12 | 0.00 | 0.02 |
| | G | 99.89 | 0.00 | 0.00 | 80.32 | 0.01 | 0.19 | 0.00 | 0.00 | 0.00 | 0.01 | 99.95 | 0.00 | 0.00 | 0.00 | 0.01 | 99.91 | 0.00 | 0.10 | 99.95 | 0.00 | 0.02 | 99.94 | 0.00 |
| | T | 0.00 | 99.95 | 99.94 | 0.04 | 99.96 | 1.28 | 99.89 | 0.04 | 0.02 | 99.87 | 0.00 | 99.96 | 99.95 | 0.02 | 99.79 | 0.01 | 0.06 | 0.02 | 0.02 | 0.03 | 99.84 | 0.01 | 99.96 |

**Fig. S22**. Off-target activities of AAV9-iNG. **a**, Sequences of on-target and two off-target sites. The target base A (green) is located at position 4 of the gRNA for all three sites. The different nucleotides are highlighted in red and the PAM sequences are in blue. **b**,

The nucleotide frequency at the on-target site of the four *mdx*$^{4cv}$ mice at 10 months after treatment with AAV9-iNG. The desired edit at $A^4$ is highlighted in green and the bystander $C^6$ edit in red.

**Table S1**. List of ABE variants engineered in this study.

| Name | Description |
|---|---|
| **ABE-NG** | ABEmax with SpCas9-NG mutations R1335V/L1111R/D1135V/G1218R/E1219F/A1322R/T1337R |
| **ABE-NGA** | ABE-NG with R1335Q mutation |
| **ABE-NGC** | ABE-NG with R1335E mutation |
| **ABE-NG-loop** | ABE-NG with the loop sequence from ScCas9 (amino acids 367-376) inserted |
| **ABE-NGX** | ABE-NG with A262T/R324L/S409I/E480K/E543D/M694I mutations |
| **ABE-NGX-NGA** | ABE-NGX with R1335Q |
| **ABE-NGX-NGC** | ABE-NGX with R1335E |
| **ABE-NGX-loop** | ABE-NGX with the loop sequence from ScCas9 (amino acids 367-376) inserted |
| **ABEmaxSc** | ABEmax with SpCas9 nickase replaced with ScCas9 nickase |
| **ABE-NGm** | ABE-NG with the dimeric TadA-TadA* replaced with monomeric TadA* containing two additional mutations A56G and V82G |
| **iABE-NG** | ABE-NGA with the dimeric TadA-TadA* replaced with monomeric TadA* containing two additional mutations A56G and V82G |

**Table S2**. List of primers used for NGS in this study.

| Name | Sequence |
|---|---|
| **Mdx4cv-E52-F** | ACACTCTTTCCCTACACGACGCTCTTCCGATCTGAACTCATTACTGCTGCCCAGA |
| **Mdx4cv-E53-R** | GTGACTGGAGTTCAGACGTGTGCTCTTCCGATCGACCTGTTCGGCTTCTTCCTTA |
| **Mdx4cv-i52-F** | ACACTCTTTCCCTACACGACGCTCTTCCGATCTAAATTTCCACTGTCTTCTCTTGAGT |
| **Mdx4cv-i53-R** | GTGACTGGAGTTCAGACGTGTGCTCTTCCGATCGCTTGCCTCTGACCTGTCCTAT |
| **mChr16OT-F** | ACACTCTTTCCCTACACGACGCTCTTCCGATCTGTGACTAGGGGCAAAGCAAGAT |
| **mChr16OT-R** | GTGACTGGAGTTCAGACGTGTGCTCTTCCGATCCTTCCAAACTTTCTGCCCATTC |
| **mChr1OT-F** | ACACTCTTTCCCTACACGACGCTCTTCCGATCTAACACAGCGTGCTCTTTCCTTAC |
| **mChr1OT-R** | GTGACTGGAGTTCAGACGTGTGCTCTTCCGATCGTTCAGAAGAACATCCCGTTGAC |
| **NGS-final-F** | AATGATACGGCGACCACCGAGATCTACACTCTTTCCCTACACGAC |
| **NGS-final-R1** | CAAGCAGAAGACGGCATACGAGATCTTGTAGTGACTGGAGTTCAGACGT |
| **NGS-final-R2** | CAAGCAGAAGACGGCATACGAGATCAGATCGTGACTGGAGTTCAGACGT |
| **NGS-final-R3** | CAAGCAGAAGACGGCATACGAGATCCGTCCGTGACTGGAGTTCAGACGT |
| **NGS-final-R4** | CAAGCAGAAGACGGCATACGAGATATGTCAGTGACTGGAGTTCAGACGT |
| **NGS-final-R5** | CAAGCAGAAGACGGCATACGAGAT GTCCGC GTGACTGGAGTTCAGACGT |
| **NGS-final-R6** | CAAGCAGAAGACGGCATACGAGAT TTAGGC GTGACTGGAGTTCAGACGT |
| **NGS-final-R7** | CAAGCAGAAGACGGCATACGAGAT CGATGT GTGACTGGAGTTCAGACGT |
| **NGS-final-R8** | CAAGCAGAAGACGGCATACGAGAT TGACCA GTGACTGGAGTTCAGACGT |
| **NGS-final-R9** | CAAGCAGAAGACGGCATACGAGAT AGTCAA GTGACTGGAGTTCAGACGT |
| **NGS-final-R10** | CAAGCAGAAGACGGCATACGAGAT AGTTCC GTGACTGGAGTTCAGACGT |
| **NGS-final-R11** | CAAGCAGAAGACGGCATACGAGAT GATCAG GTGACTGGAGTTCAGACGT |
| **NGS-final-R12** | CAAGCAGAAGACGGCATACGAGAT ACAGTG GTGACTGGAGTTCAGACGT |
| **NGS-final-R13** | CAAGCAGAAGACGGCATACGAGAT TATACT GTGACTGGAGTTCAGACGT |
| **NGS-final-R14** | CAAGCAGAAGACGGCATACGAGAT CAACAA GTGACTGGAGTTCAGACGT |
| **NGS-final-R15** | CAAGCAGAAGACGGCATACGAGAT GTTGTT GTGACTGGAGTTCAGACGT |
| **NGS-final-R16** | CAAGCAGAAGACGGCATACGAGAT TCGGTT GTGACTGGAGTTCAGACGT |
| **NGS-final-R17** | CAAGCAGAAGACGGCATACGAGAT AGTATT GTGACTGGAGTTCAGACGT |
| **NGS-final-R18** | CAAGCAGAAGACGGCATACGAGAT TCTTGT GTGACTGGAGTTCAGACGT |
